# Methods for precise targeting and modulation of the axon initial segment

**DOI:** 10.64898/2026.11.24.768858

**Authors:** Keyu Chen, Yubing Li, Fang Liu, Ming Shi, Yaojun He, Qiufeng Teng, Weitao Chen, Wenqiang Zhang, Hongze Mao, Mingjie Zhang

## Abstract

The axon initial segment (AIS) is fundamental to action potential initiation, neuronal polarity maintenance, and plasticity. Despite its functional importance, AIS remains understudied partially due to the lack of molecular tools with high subcellular specificity. By engineering an ultra-high affinity (K_d_ ∼0.37 nM) small binder (16 kDa) that exclusively targets 480-kDa ankyrin-G, the master scaffold of AIS, and integrating neuronal type-specific promoters with feedback-regulated expression elements, we developed an efficient, portable method for delivering tightly controlled amounts of proteins or organelles to the AIS with unprecedented specificity, enabling selective, tunable, and near-endogenous-level AIS targeting in defined neuronal populations both *in vitro* and *in vivo*. Coupled with a toolkit of AIS modulators that we characterized, this strategy achieves directed manipulation of AIS structure, real-time monitoring of AIS plasticity dynamics and fast degradation of the entire AIS compartment, in desired neuronal types. Thus, our method provides new opportunities for dissecting AIS physiology and pathology, facilitating the development of therapeutics for AIS-linked brain disorders.

**Significance statement:** The axon initial segment (AIS) is a master regulator of neuronal excitability and polarity, yet its study has been hindered by a lack of tools for specific, non-perturbative manipulations. Here, we introduce EATT, a compact, modular, and cell-type-specific method that achieves faithful, near-endogenous-level AIS targeting of proteins via an ultra-high affinity ankyrin-G binder. EATT enables long-term real-time imaging of AIS plasticity dynamics, directed modulation of AIS structure, and rapid inducible AIS degradation in defined neuronal populations both in vitro and in vivo. This toolkit bridges a critical methodological gap, opening new avenues for dissecting AIS physiology and pathology, and accelerating therapeutic development for AIS-linked brain diseases.

## Introduction

Compartmentalization is a foundational principle of cellular architecture, as it allows distinct subcellular domains to execute specialized functions. Elucidating molecular mechanisms that govern compartment-specific structure and function is a major research field of cell biology. Effective tools and methods that can specifically monitor and target molecular components in individual subcellular compartment are drivers for the advancements of various fields.

The axon initial segment (AIS) of neurons is a prime example of a subcellular compartment that is highly specialized and paramount to the physiological functions of neurons ^1,2^. Located at the soma-axon junction and with a size of ∼1 μm in diameter and 30∼60 μm in length, the AIS maintains soma/dendrite-axon polarity by acting as a selective barrier that controls the entry of proteins, vesicles, and organelles into the axon, thereby supporting neuronal polarization and the initiation and propagation of action potentials ^3,4^. This gating function of the AIS is dynamic and tightly regulated ^5–8^, but with understudied molecular mechanisms.

Furthermore, the AIS is crucial for neuronal function by integrating electrical signals and initiating action potentials across neuronal types ^9,10^. It also receives inhibitory inputs from chandelier interneurons via axo-axonic GABAergic synapses, which are exclusively formed at the AIS to inhibit target neuron firing and modulate spike timing/excitation-inhibition balance ^11,12^. Thus, the AIS is a pivotal subcellular compartment for modulating neuronal output.

As the critical gatekeeper of neuronal output, the AIS dynamically adjusts its length and subcellular position along the proximal axon in response to altered neuronal activities, thereby serving as a homeostatic regulator that maintains the excitation-inhibition (E/I) balance of neural circuits ^13–17^. Dysregulation of the E/I balance is increasingly recognized as a pathological hallmark of diverse neurological and psychiatric disorders ^18,19^, with accumulating evidence linking impaired AIS plasticity to conditions including epilepsy, Alzheimer’s disease, bipolar disorders, schizophrenia, neuroinflammatory and demyelinating pathologies ^20–24^. A recent study also reveals that AIS plasticity is not only vital for homeostatic adaptation but also a hallmark of memory formation ^25^. Thus, deciphering the principles underlying the AIS plasticity regulation will deepen our comprehension of how AIS dysfunction contributes to disease pathologies and facilitates the development of therapeutics for AIS malfunction-related diseases.

However, a large knowledge gap exists between the functions of AIS and the underpinning molecular mechanisms. Except for several AIS structural proteins, such as the 480-kDa giant ankyrin-G (gAnkG) and βIV-spectrin ^1,2^, the majority of AIS proteins (various AIS regulatory proteins in particular) also exist in other subcellular regions of neurons. Because the AIS constitutes only a tiny fraction of the total neuronal volume, only a very minor portion of each regulatory protein is localized to the AIS. Consequently, methods that target these AIS regulatory proteins inevitably also alter their functions in other subcellular locations. Additionally, specific neuronal cell type-selective regulation strategies for the AIS are also highly desired but currently lacking.

To overcome these challenges, we engineered a novel AIS-targeting platform by identifying and optimizing a small, stable, selective, and potent gAnkG-binding protein. By combining neuron type-specific promoters and feedback-regulated expression of this gAnkG-binding protein, we achieved robust and faithful targeting of any desired proteins or even organelles to the AIS in living neurons, allowing specific monitoring of the AIS dynamics at single-neuron resolution, dictating structural changes of the AIS, and altering AIS-resident molecular components in living neurons. In summary, our approach provides a transformative tool for exploring AIS physiology under native conditions and holds promise for interrogating neurological disorders associated with AIS dysfunction.

## Results

### Rational design of a genetically encodable AIS-targeting tool

Targeting specific subcellular compartments in living cells requires genetically encodable binders that possess high affinity for their target proteins and interact with them exclusively within the intended compartment. Uncontrolled ectopic expression of such binders often encounters multiple problems, such as unstable expression and batch-to-batch variation. Insufficient expression leads to poor target occupation, whereas overexpression causes the binder to overflow to locations other than the desired compartment. We therefore incorporated a feedback-controlled expression system ^26^ into our AIS-targeting tool (Figure 1a). The construct includes a gAnkG binding domain (ABD) for specific AIS targeting, a CCR5 Zinc Finger (ZnF) domain to recruit the Krüppel-associated box (Krab) domain to the promoter for transcriptional repression ^26–28^, and a woodchuck hepatitis virus post-transcriptional regulatory element (WPRE) for mRNA stabilization. This design ensures that, upon saturation of the AIS binding sites, excess tool protein diffuses to the nucleus and suppresses its own transcription via the CCR5-Krab elements (Figure 1b).

**Figure 1:**
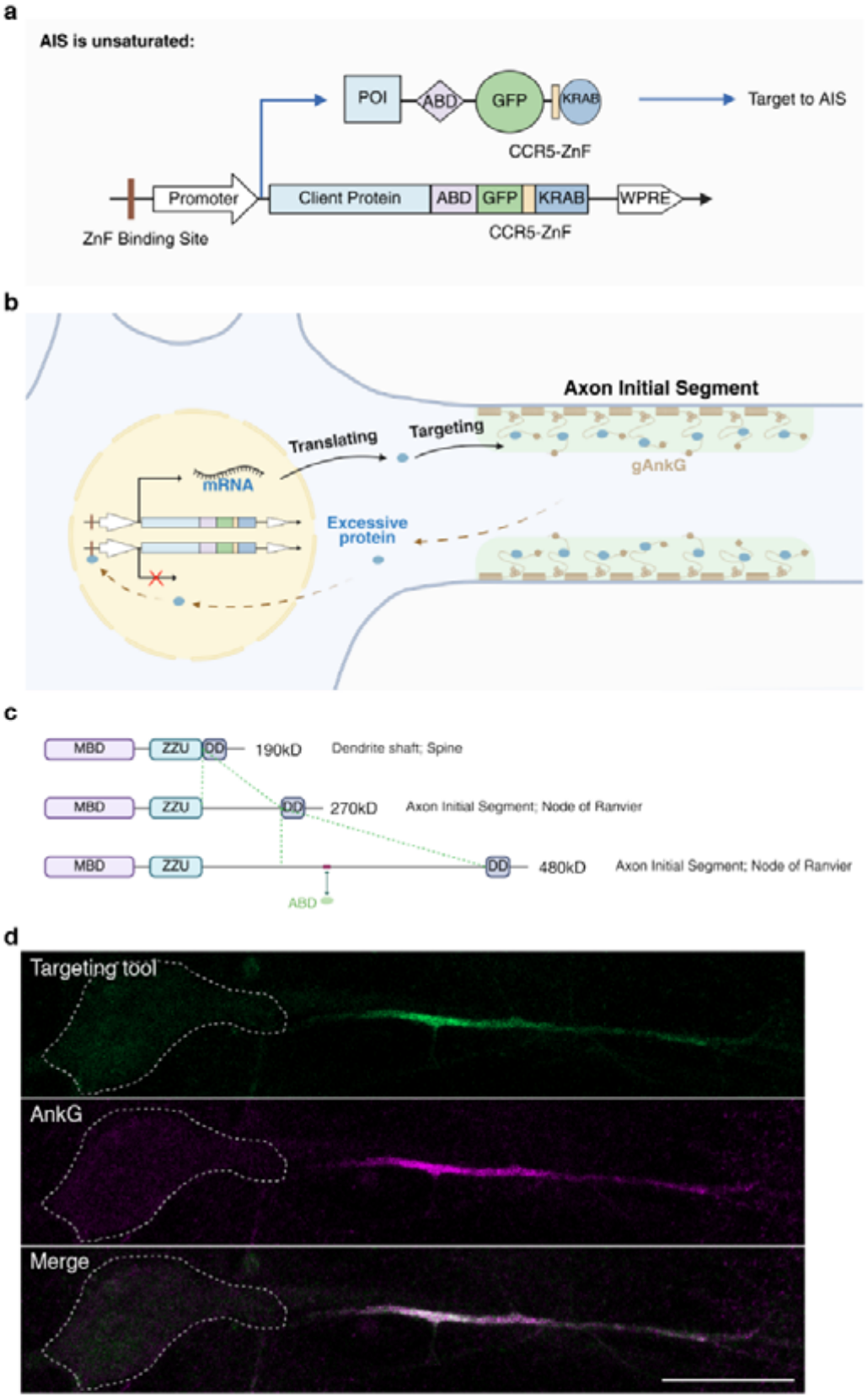
Design of a genetically encodable AIS targeting tool. **a:** Schematic diagram of the AIS targeting tool construct. POI: protein of interest; ABD: AnkG-binding domain; CCR5-ZnF: CCR5-like zinc finger; WPRE: Woodchuck hepatitis virus posttranscriptional regulatory element; ZBS: ZnF-binding site, a cis-regulatory element interacting with CCR5-ZnF. b: Illustration of the principle of AIS targeting and feedback-controlled expression. **c:** Domain organization of major AnkG isoforms (encoded by the *Ank3* gene) in neurons and their subcellular localizations. The ABD from MCF2L and its binding site on 480-kDa AnkG are indicated by a two-headed arrow. **d:** Specific labeling of the AIS in a DIV9 hippocampal neuron by the AIS targeting tool. Green: fluorescence signal from EGFP tag of the tool; magenta: immunostaining of endogenous AnkG. Scale bar: 20 μm.

Multiple splicing isoforms of AnkG are expressed in neurons (Figure 1c) ^29,30^. Among these, the 480-kDa giant Ankyrin-G (gAnkG), almost exclusively clustered at the AIS and Nodes of Ranvier (an evolutionarily homologous structure to the AIS), serves as the central organizer and is responsive to nearly all forms of AIS plasticity ^2,31,32^. These properties make gAnkG an ideal anchoring point for AIS-targeting tools. Furthermore, gAnkG is one of the most abundant proteins decorating the entire AIS. Thus, delivering proteins at levels sufficient to saturate gAnkG should achieve concentrations high enough for most experimental purposes.

We evaluated several known gAnkG binders with small molecular sizes and nanomolar binding affinities, including GABARAP (K_d_= 2.6 ± 0.7 nM; with its C-terminal lipidation sequence removed) ^33^ and scFv N106-20, a single-chain variable fragment derived from a gAnkG monoclonal antibody ^34^. In addition, our previous yeast two-hybrid screening using a fragment (residues 2329-2691) of the 480-kDa isoform-specific sequence (the giant insertion) as bait identified MCF2L, a multi-domain RhoGEF family member containing an N-terminal Sec14 domain, three spectrin repeats, DH, PH, and SH3 domains ^35^, as a novel gAnkG binder that selectively interacts with gAnkG rather than other isoforms (Figure 1c) ^36^; the interaction is mediated specifically by the first spectrin repeat (Spec1) of MCF2L and exhibits high affinity (K_d_ = 8.78 ± 1.46 nM, Figure 2d). Detailed identification and characterization of this interaction have been reported by Li et al. ^36^. GABARAP accumulated in the nucleus, whereas the gAnkG_scFv showed diffuse cytosolic localization, failing to specifically label the AIS (Figure S1a, S1b). In contrast, MCF2L Spec1 effectively clustered at the AIS (Figure 1d). A single point mutation in this domain disrupted its gAnkG binding and abolished the AIS targeting (Figure S1c). The MCF2L Spec1 is small in size (∼16 kDa), highly soluble, stable, freely diffusible, and has a short binding site that only exists in the 480-kDa gAnkG (Figure 1c). We designated this construct EATT1.0 (Endogenous-level-AIS-Targeting Tool, version 1.0).

**Figure 2:**
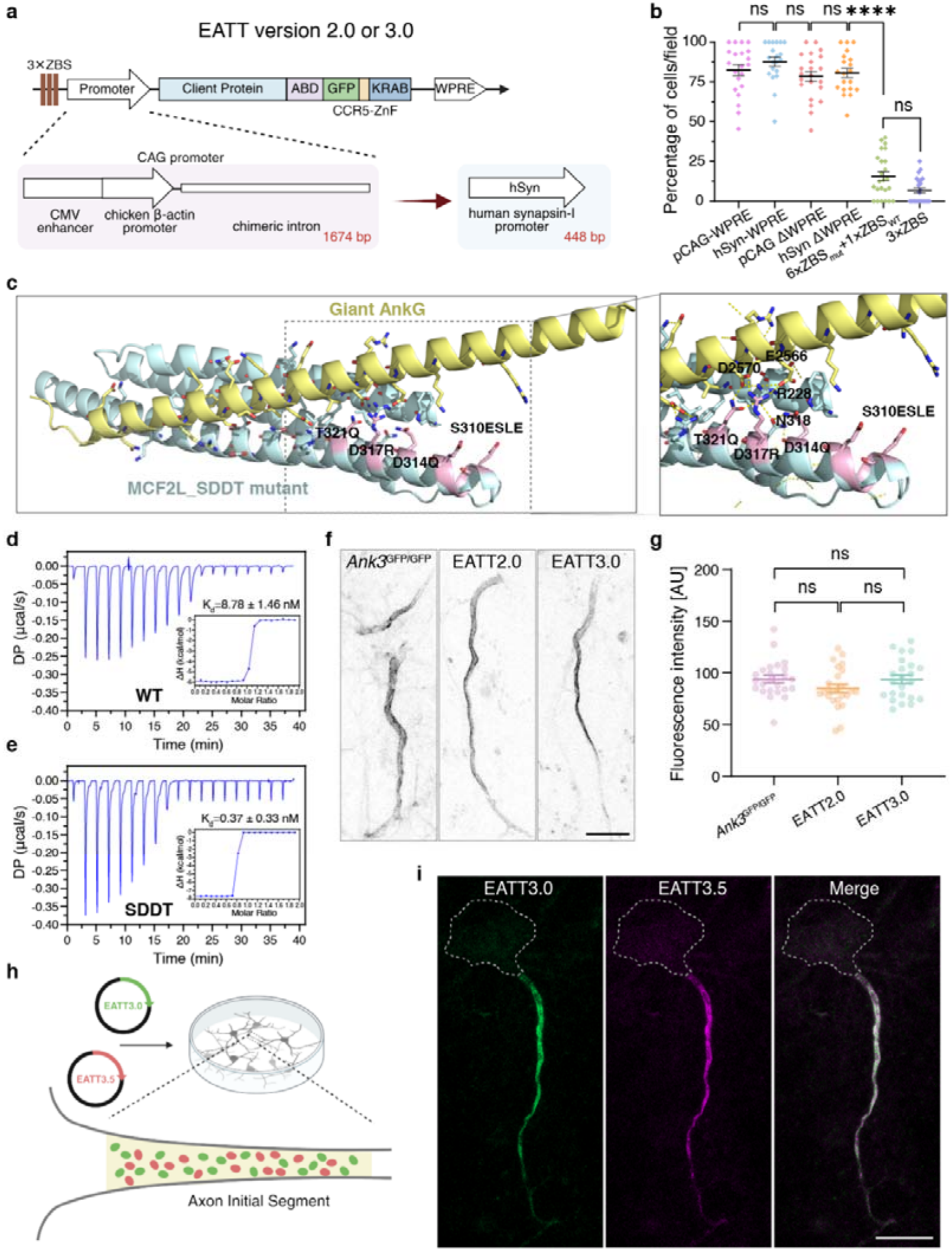
Optimization and characterization of the AIS targeting tool. **a:** Schematic showing promoter substitution from CAG to human Synapsin-1 (hSyn). **b:** Quantification of the ratio of neurons with nuclear ABD-GFP accumulation. Signals are from GFP fluorescence without immunostaining. Nuclear accumulation was defined as nuclear GFP intensity >50% of AIS GFP intensity. Data are from random optical fields, mean ± SEM; *n* > 20 fields per group; Kruskal-Wallis test followed by Dunn’s multiple-comparison test; ns, P>0.05; ****P < 0.0001. **c:** AlphaFold III-predicted complex structure of the MCF2L ABD SDDT mutant (light cyan) and the gAnkG fragment (pale yellow). Mutated residues (light pink) and key interaction residues (sticks) are shown. The boxed region is enlarged. **d, e:** Isothermal titration calorimetry (ITC) measurements of binding affinity between the gAnkG fragment and MCF2L ABD WT (**d**, *K_d_* = 8.78 ± 1.46 nM) or SDDT mutant (**e**, *K_d_*= 0.37 ± 0.33 nM). **f:** Hippocampal neurons from AnkG-GFP knock-in or WT C57BL/6J mice transfected with EATT2.0 or EATT3.0. Imaging used identical confocal settings without immunostaining and are displayed in inverted gray scale. **g:** AIS fluorescence intensity from (**f**). Student’s *t* test, *n* > 23 neurons per group, mean ± SEM. **h:** Schematic of orthogonal AIS targeting tools carrying green or red fluorescent proteins respectively. Co-transfection of them leads to simultaneous targeting of both proteins to the AIS. **I:** Images showing simultaneous and specific AIS labeling by orthogonal EATTs. Scale bar: 20 μm.

### Optimization and characterization of the AIS-targeting tool

Although EATT1.0 successfully labeled the AIS, a substantial portion of transfected cells exhibited bright nuclear signals (Figure S1d). This phenomenon, commonly observed in other CCR5-Krab-based systems ^26,37–39^, is possibly caused by pulsatile expression waves. To stabilize expression, we employed several strategies, either individually or in combination: (1) removing the WPRE element to reduce peak expression levels; (2) substituting the original promoter with a more stable one to lower the peak expression and enhance the long-term expression stability; (3) increasing the copy number of the ZnF binding sequence (ZBS, a cis-regulatory element) to enhance the avidity for the ZnF domain and thus strengthen the feedback regulation.

Replacing the original CAG promoter in EATT1.0 with the human Synapsin-1 (hSyn) promoter did not significantly alter its expression pattern (Figure 2a, 2b), consistent with reports using these promoters for other purposes ^40,41^. Given its smaller size (advantageous for viral packaging) and its neuron-specific expression, we adopted the hSyn promoter for our subsequent designs (Figure 2a). Further removal of the WPRE element slightly decreased nuclear accumulation of the expressed AIS targeting proteins (Figure 2b).

Inspired by a discovery showing natural existence of short tandem cis-element repeats in tuning gene expression ^42^, we introduced multiple copies of either the wild-type ZBS (ZBS_WT_) or mutant ZBS (ZBS_mut_, which has a weaker binding affinity for ZnF) into the hSyn promoter-containing construct to tune the stringency of the feedback control. Remarkably, incorporating six copies of ZBS_mut_ plus one copy of ZBS_WT_ (i.e., 6× ZBS_mut_ +1× ZBS_WT_) significantly stabilized expression and reduced nuclear accumulation of ABD-GFP (Figure 2b). Tripling the ZBS_WT_ copy number (i.e., 3× ZBS_WT_, Figure 2a) also suppressed leakage, with over 90% of transfected cells exhibiting no detectable nuclear GFP signal (Figure 2b). This optimized system was designated as EATT2.0 (see Figure S1e for schematics of all EATT versions).

Previously, we solved the crystal structure of the ABD from MCF2L in complex with its binding segment from gAnkG ^36^. Aided by AlphaFold III prediction and based on the complex structure, we designed an ABD mutant with an enhanced affinity to gAnkG. In this mutant, Ser310 was substituted with a four-residue insertion (“Glu-Leu-Ser-Glu”, ELSE) to stabilize the α3-helix of the spectrin repeat; and three substitutions (D317R, D314Q and T321Q) were introduced to enhance polar interactions at the ABD-gAnkG interface (Figure 2c). This ABD mutant, named SDDT, remained a stable monomer in solution (Figure S1f) and exhibited an approximately 24-fold increase in gAnkG binding affinity compared to the wild-type ABD (Figure 2d, 2e). The corresponding construct, EATT3.0 (Figure S1e), also achieved near-exclusive AIS localization (Figure 2f, S1g).

To verify that ABD-GFP expression levels approximate those of endogenous gAnkG, we generated AnkG-GFP knock-in mice with a C-terminal EGFP tag fused to the AnkG proteins (Figure S1h-S1j). Primary hippocampal neurons from these mice exhibited clear AIS enrichment of AnkG-GFP in both live and fixed conditions without immunostaining. Comparison of these signals with GFP signals from EATT2.0- or EATT3.0-transfected neurons confirmed that ABD-GFP expression was comparable to the endogenous AnkG levels at the AIS (Figure 2f, 2g). AIS labeling was detectable as early as 6 hours post-transfection (Figure S2a-b) and was maintained effectively for at least 20 days (Figure S2c-d). Notably, although the fluorescence intensities of EATT2.0 and EATT3.0 constructs showed no significant difference once stable expression was achieved (Figure 2g), EATT3.0 was significantly brighter than EATT2.0 during the early expression stages (Figure S2e), consistent with more efficient initial recruitment of EATT3.0 due to its higher gAnkG binding affinity.

We further developed an orthogonal AIS-targeting tool by replacing the CCR5-Krab and the ZBS cassette with IL2R-Krab and its corresponding ZBS, swapping the hSyn promoter for the EF1α promoter, and substituting GFP with mCherry ^26^ (Figure 2h). This variant, designated EATT3.5 (Figure S1e), enabled simultaneous and perfectly overlapped labeling of the AIS by ABD-GFP and ABD-mCherry (Figure 2i). These orthogonal AIS-targeting tools enable compartment-restricted co-localization of multiple proteins, providing another layer of spatial-specificity control when combined with split-protein systems (e.g., split-BioID), critical for applications demanding stringent subcellular confinement of proteins.

### Potent and specific AIS-targeting ability of the EATT system

We then assessed whether EATT application perturbs AIS structure. Transfection of EATT2.0 into cultured hippocampal neurons revealed no structural alterations in the AIS, as marked by gAnkG immunostaining, compared with non-transfected cells (Fig. 3a, 3b). Similar results were observed for EATT3.0 (Fig. S3a). In addition, staining for pan-NaV, Neurofascin and βIV-spectrin in EATT2.0-transfected neurons showed no alteration in the distribution or intensity of these AIS hallmark proteins (Figure 3c-g and larger optic views in S3b, S3c). This finding was further validated *in vivo* by expressing ABD-GFP in mice brain using AAV-packaged EATT2.0, which did not affect the expression patterns of gAnkG and βIV-spectrin (Figure S3d-S3g). Thus, the expression of EATT does not compromise the AIS integrity both *in vitro* and *in vivo*.

**Figure 3:**
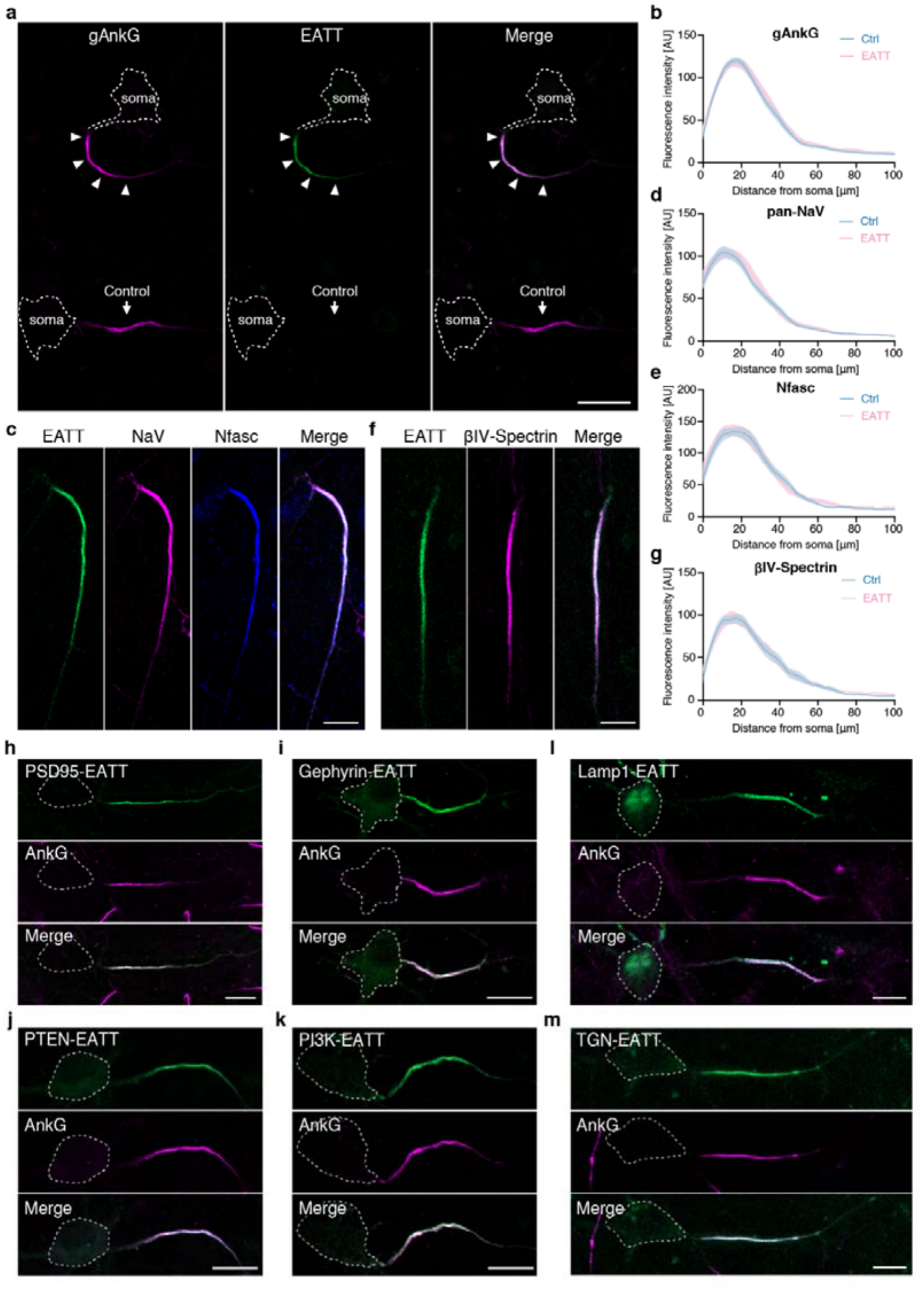
Efficient and specific AIS targeting abilities of the EATT system. **a:** Representative images of AnkG immunostaining showing no morphological differences in the AIS between tool-expressing and untransfected neurons. Scale bar: 20 μm. **b:** Profile plot of gAnkG immunostaining signal along the axon. The raw intensity curves are smoothened for better presentation. Solid lines indicate mean values; shaded bands represent ± SEM. Ctrl, n=34; EATT, n=33. **c-g:** AIS images and intensity profiles of Pan-NaV (**c, d**), Neurofascin (**c, e**) or βIV-spectrin (**f, g**) immunostaining in EATT-expressing neurons, indicating no differences compared to untransfected neurons. Scale bar: 10 μm. Solid lines, mean; shaded areas, mean ± SEM. n=16 (EATT) and 26 (Ctrl) in **d**; n=18 (EATT) and 27 (Ctrl) in **e**; n=21 (EATT) and 20 (Ctrl) in **g**. **h-k:** Images of hippocampal neuron AIS targeted by PSD95-EATT (**h**), Gephyrin-EATT (**i**), PTEN-EATT (**j**), and PI3K-EATT (**k**), respectively. Endogenous AnkG was immunostained with a pan AnkG antibody. Somata are outlined with dashed circles. Scale bar: 10 μm. **l-m:** Images of lysosomes (**l**) or trans-Golgi vesicles (**m**) targeted to the AIS by EATT. Scale bar: 10 μm.

Large scaffold proteins, due to their robust and extensive interaction networks at specific subcellular compartments, are typically resistant to ectopic redistribution. To test whether EATT2.0 can enrich such proteins at the AIS, we fused the C-terminus of ABD-GFP to several postsynaptic scaffold proteins, PSD95 (80.5 kDa), CNKSR2 (117.4 kDa), and Gephyrin (83.3 kDa), which normally localize to excitatory (PSD95, CNKSR2) or inhibitory (Gephyrin) postsynaptic densities. Remarkably, expression of these fusion proteins using the EATT2.0 system led to their precise and pronounced enrichment at the AIS as evidenced by their perfect colocalization with the endogenous gAnkG (Figure 3h, 3i and S4a; see the original localization of these proteins in Figure S4b), underscoring the robust capability of our tool in specifically recruiting and concentrating even large scaffold proteins to the AIS.

We next evaluated whether the EATT system can target cytosolic enzymes specifically to the AIS, a strategy that is important for investigating their regulatory roles, given that most of these enzymes function in multiple cellular compartments. We demonstrated that PTEN and PI3K, two key enzymes involved in phosphatidylinositol phosphate metabolism, could also be specifically targeted and clustered at the AIS when expressed as ABD-GFP fusions (Figure 3j, 3k and S4b). This approach provides opportunities for investigating the roles of specific phosphatidylinositol phosphate lipids in the AIS.

The distribution of organelles within neurons is uneven and critical for their cellular functions ^43,44^. For instance, lysosomes are predominantly confined to the soma, with only a small fraction entering axons. By fusing Lamp1, a late endosome/lysosome integral membrane protein, to the N-terminus of ABD-GFP, we observed a striking redistribution of a portion of lysosomes to the AIS with excellent colocalization with gAnkG, and as expected most lysosomes remained in the soma (Figure 3l, S4b). This tool can be used to study functional impacts of enhanced lysosome activities at the AIS. Conversely, the EATT system also enables specific interference with AIS-localized lysosomes by fusing lysosomal regulatory factors to ABD-GFP. On the other hand, certain somatodendritic organelles and vesicles, are typically excluded from the axons by the AIS. We explored whether our EATT system could override this gating restriction by actively driving and clustering of these organelles to the AIS. The Golgi apparatus, for example, is known to be excluded from AIS ^45^. Fusing TGN38, a resident protein of the trans-Golgi network, to ABD-GFP in the EATT2.0 system, resulted in clustering of TGN38-labeled vesicles at the AIS (Figure 3m, S4b). Additionally, our system proved capable of enriching even very large organelles, such as mitochondria, at the AIS, as demonstrated by fusing the outer mitochondrial membrane protein mitofusin 1 (MFN1) to ABD-GFP (Figure S4b, S4c).

### Cell-type selective AIS-targeting by the EATT system

Tools capable of differentially labeling or targeting to the AIS in specific neuronal populations are highly desirable. While transgenic mouse models can achieve this, such efforts are often time-consuming and costly. We therefore explored the adaptability of our EATT system by coupling it with neuron-type-specific promoters to label distinct neuronal populations in cultures. We replaced the hSyn promoter in EATT2.0 with either the CamkIIα or the hDlx promoters, directing AAV-mediated expression of ABD-GFP to glutamatergic excitatory or GABAergic inhibitory neurons, respectively, while keeping all other elements unchanged (Figure 4a). As expected, ABD-GFP faithfully clustered at the AIS in the corresponding neuronal types (Figure 4b-4e). Immunostaining against CaMKII confirmed that about 97.5% of neurons expressing ABD-GFP under the CamkIIα promoter were correctly identified as glutamatergic (Figure 4c). Similarly, about 95.8% of neurons expressing ABD-GFP driven by the hDlx promoter were co-labeled with the inhibitory marker Gad67 (Figure 4e). We further validated this selective targeting *in vivo* using AAV-mediated delivery of the promoter-specific EATT2.0 systems in the mouse cortex, which similarly enabled selective AIS labeling in glutamatergic or GABAergic neurons, respectively (Figure 4f, 4g). These findings demonstrate the high versatility and specificity of our EATT platform for cell type-selective AIS targeting both *in vitro* and *in vivo*.

**Figure 4:**
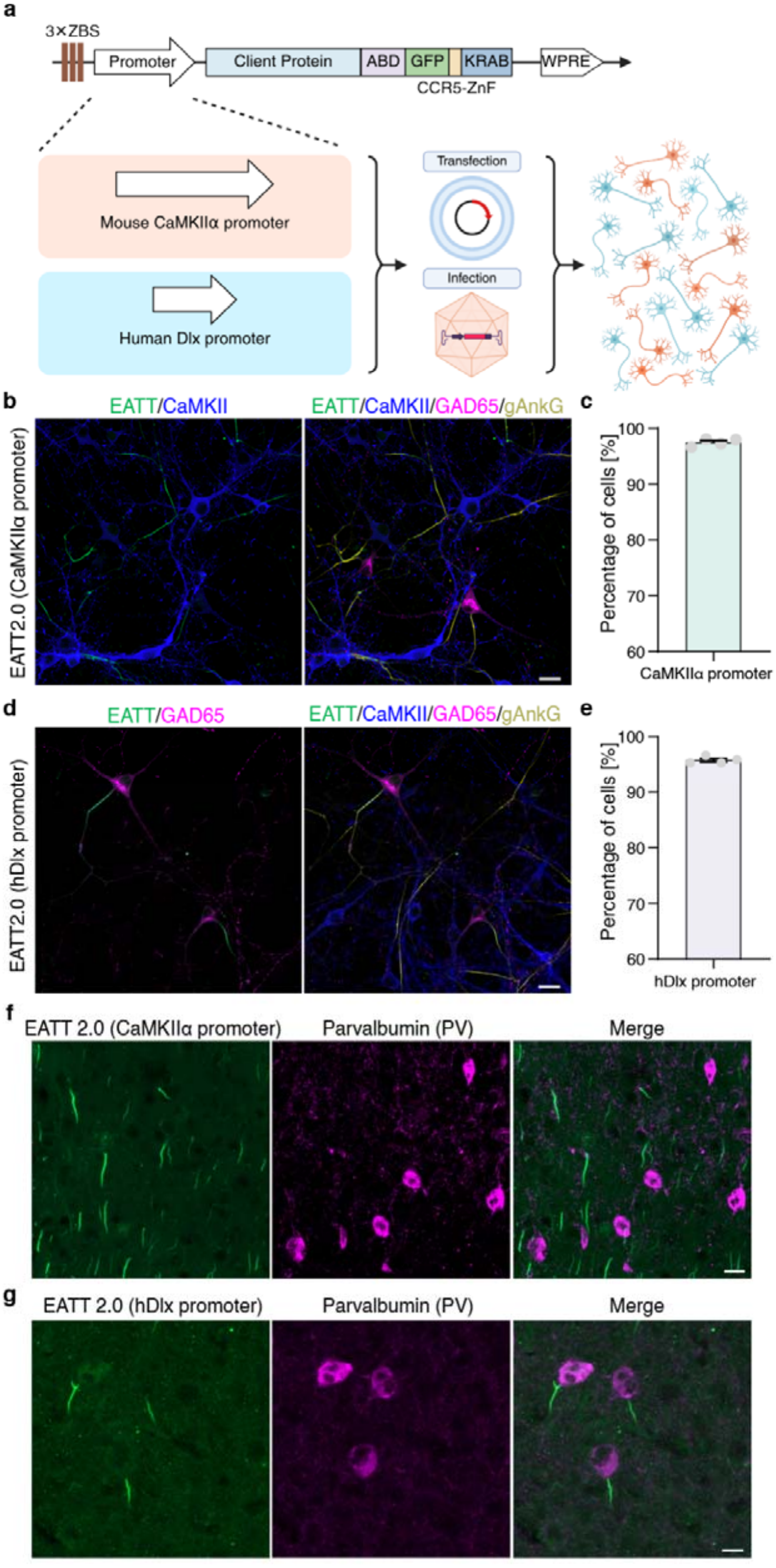
Cell type-specific AIS targeting by the EATT system. **a:** Schematic of the modular EATT design. The promoter module is interchangeable to enable cell type-specific targeting via DNA transfection or viral delivery. **b:** AAV-delivered EATT (green) driven by the Camk2a promoter selectively labels glutamatergic neurons (blue, anti-Camk2a); GABAergic neurons (red, anti-Gad65) and all AISs are immunostained with anti-gAnkG (yellow) in right panel. Scale bar, 20 μm. **c:** Percentage of EATT (Camk2a promoter)-labeled neurons positive for anti-Camk2a. Each data point represents one dish; fields were imaged randomly without overlapping until 150 cells were counted or all fields captured. Mean ± SEM. **d:** As in b, but with EATT driven by the hDlx promoter, selectively labeling GABAergic neurons (green). Glutamatergic neurons (blue) and all AISs (yellow) are labeled in right panel. Scale bar, 20 μm. **e:** Percentage of EATT (hDlx promoter)-labeled neurons positive for anti-Gad65; imaging as in **c**. **f:** Labeling of excitatory neuron AIS in adult mouse motor cortex 30 days after injection with AAV9-packaged EATT (Camk2α promoter, green) shows selective labeling of excitatory neurons. GABAergic interneurons were immunostained for Parvalbumin (PV; red). The EATT signal is notably absent from the AIS of PV-positive neurons. Scale bar: 20 μm. **g:** Labeling of inhibitory neuron AIS in mouse motor cortex 30 days after injection with AAV9-packaged EATT (hDlx promoter, green) shows expression in GABAergic neurons instead of excitatory neurons. PV-positive interneurons are immunostained (red). Scale bar: 20 μm.

### Directed manipulation of AIS structure using the EATT system and newly characterized modulators

The AIS is a stable but plastic structure, whose dimensions and molecular composition can adapt in response to neuronal activities ^13,14^. Notably, the structural properties of the AIS, such as its size and location relative to soma, determine the activity pattern of neurons ^17^. Although AIS plasticity has been documented under diverse physiological and pathological conditions ^2,46–52^, selective manipulation of specific AIS architectural features remains challenging due to the lack of known remodeling factors and approaches to exclusively deliver them to the AIS without inducing off-target effects in other neuronal domains. To address this, we employed the EATT system to deliver various candidate protein effectors to the AIS, evaluating their AIS-targeting specificity and structural remodeling capacity. This approach yielded a versatile toolkit conferring multi-dimensional control over the AIS structure, including strengthening or weakening its structural integrity, modulating its distance from the soma, altering its length or diameter, and even inducing AIS branching or the formation of multiple AIS/axons in a single neuron (Figure 5a).

**Figure 5:**
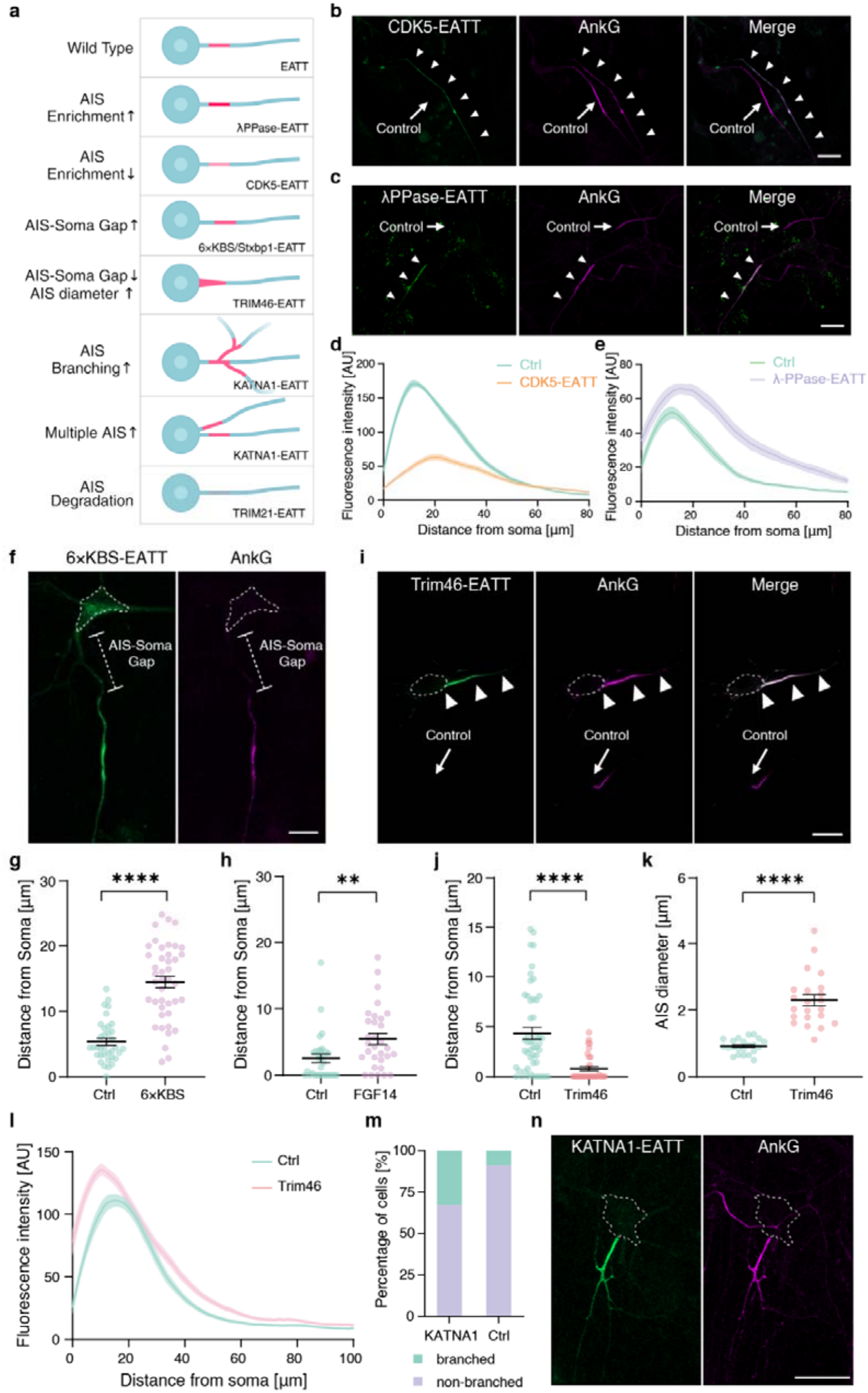
Modulation of the AIS structure using the EATT system and newly characterized modulators. **a:** A summary scheme of the AIS manipulation/modulation examples. The AIS are indicated in red or pink. **b:** CDK5-EATT (Green) decreases AnkG (magenta) intensity at the AIS. The CDK5-targeted AIS is indicated by triangles; the control AIS is indicated by an arrow. Scale bar: 20 μm. **c:** λPPase-EATT (green) increases AnkG (magenta) intensity at the AIS. The targeted and control AISs are indicated as in (**b**). Scale bar, 20 μm. **d-e:** gAnkG immunofluorescence profiles along axons in CDK5-targeted (**d,** n=50 for CDK5-EATT or Ctrl) or λPPase-targeted and control neurons (**e,** n=21 and 20 for λPPase-EATT and Ctrl respectively). Curves are smoothened, mean ± SEM. **f-g:** Images and quantification showing that EATT with six copies of the kinesin binding sequence (6×KBS) increases the soma-AIS gap. Scale bar, 10 μm; Mann-Whitney test, mean ± SEM. Scale bar: 10 μm. **h:** Quantification showing that an EATT carrying FGF14 significantly increases the soma-AIS gap. Mann-Whitney test, n=31, mean ± SEM. **i:** Images showing that EATT with Trim46 dramatically alters AIS morphology. The arrow indicates the control AIS. Scale bar, 20 μm. **j-k:** Quantifications showing that Trim46-targeted AIS have a larger diameter (Student’s t-test, P<0.0001) and a smaller soma-AIS gap (Mann-Whitney test), n=33 (Trim46) and 50 (Ctrl); mean ± SEM. **l:** gAnkG immunostaining profiles along axons showing higher intensity in Trim46-targeted AISs. Curves are smoothened, n=33 (Trim46) and 50 (Ctrl); mean ± SEM. **m-n:** Images (**m**) and quantification (**n**) showing that targeting KATNA1 to the AIS induces AIS branching. 26 and 30 cells counted for KATNA1 and control respectively. Scale bar: 20 μm.

CDK5 is known to function in multiple subcellular compartments, including the AIS ^53,54^. To visualize how CDK5 activation may modulate the AIS structure, we specifically expressed the constitutively active form of CDK5 (generated by fusing its minimal activation domain P16 to CDK5 via a flexible GS linker ^55^) using the EATT2.0 system in cultured hippocampal neurons. Interestingly, active CDK5 at the AIS caused a significant weakening of the AIS structure, as judged by decreased clustering and lower signal intensity of gAnkG compared to untransfected controls in the same culture (Figure 5b, 5d). Conversely, AIS-targeted expression of bacteriophage lambda phosphatase (λ-PPase), a broad-spectrum phosphatase, enhanced gAnkG clustering and strengthened the AIS (Figure 5c, 5e). These data indicate that the AIS structural integrity can be bidirectionally modulated by the phosphorylation status of its constituent proteins.

The change in the AIS-soma gap is a critical dimension of AIS plasticity. Using the EATT system, we discovered several new factors that can modulate this gap when targeted to the AIS. Molecular motors such as kinesins and myosins have been implicated in AIS structure plasticity ^56,57^. Fusing six copies of the kinesin-binding-sequence (6×KBS; each KBS is a 10-residue KIF5 light chain binding sequence from its cargo adaptor SKIP ^58^) to ABD-GFP induced a dramatic distal shift of the AIS, increasing the AIS-soma gap from ∼5 µm in controls to ∼14 µm in the targeted cells (Figure 5f, 5g). A similar distal shift was observed with Stxbp1-EATT, likely due to its synaptic vesicle localization and associated motor activity on these vesicles (Figure S5a, S5b). FGF14 is a small intracellular modulator of voltage-gated sodium channels and potassium channels ^59,60^. FGF14 fused to ADB-GFP expressed at the AIS led to a significant increase of the AIS-soma gap (Figure 5h, S5c), hinting that activation of the Nav channels can induce AIS plasticity.

Trim46 is a cytoskeleton-associated protein localized at the axon hillock, organizing microtubules at the AIS but not involved in axon specification or initial segment formation ^45,61–64^. The localizations of Trim46 and gAnkG only partially overlap, with Trim46 positioned more proximally and gAnkG more distally to the axon hillock ^45^. Interestingly, forced colocalization of Trim46 with gAnkG at the AIS using EATT led to a thicker and stronger AIS with a reduced AIS-soma gap, a morphology reminiscent of a more mature and robust AIS (Figure 5i-l), suggesting that the interplay between gAnkG and trim46 is crucial for AIS morphogenesis.

A typical neuron possesses a single AIS and axon branching normally occurs at sites distal to it. We tested whether forced targeting of several reported axon branching factors to the AIS could induce structural changes. As depicted in Figure 5m-n, forced expression of KATNA1, a microtubule-severing enzyme, at the AIS led to robust axonal branching in the AIS region. In a subset of neurons, KATNA1 expression even led to the formation of multiple axons, each with a distinct AIS (Figure S5d). Axon specification is tightly regulated to ensure that each mature neuron contains only one axon ^65,66^. Our KATNA1 data indicate that axonal branching and specification may share overlapping regulatory mechanisms.

Taken together, as a proof of concept, the above examples demonstrate that the EATT system developed here can be effectively adapted to manipulate the AIS structure, function and/or plasticity, as well as to study the action mechanisms of individual regulatory factors or their pairwise combinations using the orthogonal EATT3.5 system shown in Figure 2i. Our platform also holds promise for screening for new AIS regulatory proteins in cultured neurons.

### Real-time tracking of AIS plasticity in specific neuron types

The AIS is a major site of structural and functional plasticity in neurons, where interventions of neuronal activity *in vitro* and *in vivo* can induce changes in the AIS position and length, correlating with alterations in neuronal intrinsic excitability. However, most evidence for activity-dependent AIS changes comes from fixed samples, offering only static, population-level insights. Moreover, given the distinct AIS plasticity profiles of different neuron types ^13–15,67,68^, tools capable of labeling the AIS in specific neuronal populations would be valuable for studying AIS plasticity at single-cell resolution. Although an AnkG-GFP knock-in strategy has recently been developed for long-term and minimally perturbing AIS labeling, ^69^, its broad application is limited by the requirements of transgenic breeding. To generate more convenient tools for AIS monitoring in designated neuron types such as PV interneurons, we changed the promoter to hDlx and substituted EGFP in EATT3.0 with Baojin, a brighter and more photostable monomeric green fluorescent protein ^70^. We infected hippocampal and cortical neurons with this modified construct at DIV5 and performed live imaging at DIV13 for 26 hours or longer. Following application of 15 mM KCl to induce plasticity of the AIS ^16,71^, we captured time-dependent relocation of the AIS (Figure 6a and Movie S1). Robust and specific AIS labeling was maintained even after 48 hours of continuous imaging (Figure S6a). Quantification of the AIS-soma gap over time, normalized to the pre-treatment baseline, revealed distinct kinetic phases of KCl-induced AIS plasticity: an initial lag phase (∼4 hours) with small displacements, consistent with reported observations ^16^; a rapid shifting phase from 4 to ∼20 hours; and a subsequent plateau from 20 hours onward (Figure 6b).

**Figure 6:**
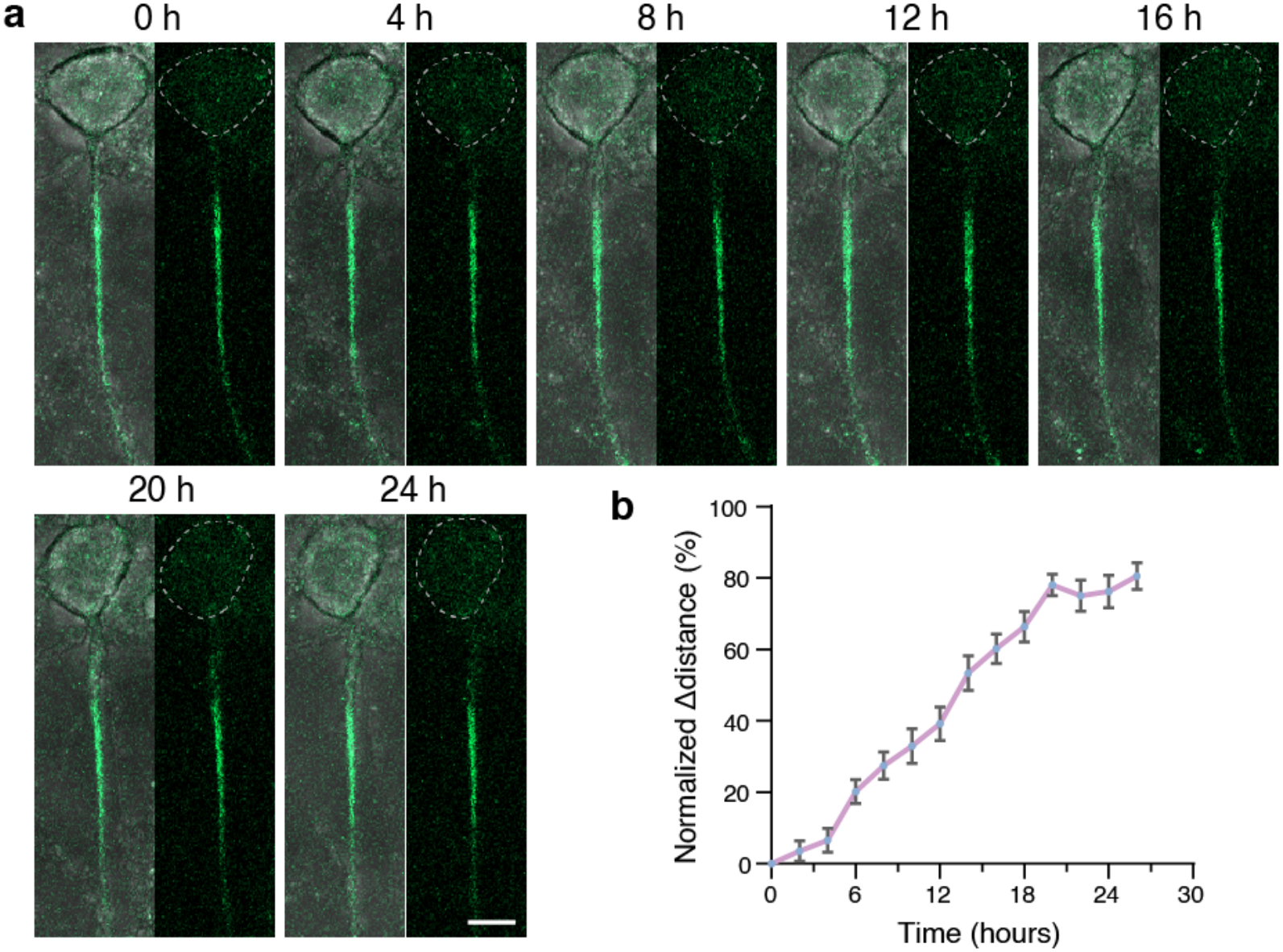
Real time monitoring of AIS plasticity in interneurons using the EATT. **a**: Snapshots of AIS positioning at indicated time points after KCl-induced plasticity, visualized by EATT-mediated Baojin expression (hDlx promoter) in live hippocampal neurons (green). The left panels are overlaid with signals from the T-PMT channel to outline the soma and AIS-soma gap. Scale bar: 10 μm. **b**: Time course of changes in the AIS-soma gap distance after addition of 15mM KCl. Δdistance was normalized such that the distance at t=0 set to 0 and the largest distance change set to 100%. n=34, mean ± SEM.

### Acute degradation of gAnkG and the AIS

Protein degradation technologies (such as Trim-Away and PROTACs) enable rapid and reversible depletion of target proteins via the ubiquitin-proteasome system and offer unique advantages over traditional genetic knockout or knockdown approaches. Genetic knock-out, which permanently eliminates a gene and often triggers compensatory mechanisms; knock-down, which usually results in only partial reduction of protein and may require several days to take effect, limitations that are particularly pronounce for scaffold proteins within exceptionally stable subcellular architectures such as the AIS ^2^. Conventional genetic ablation or knockdown of gAnkG disrupts AIS structure and leads to the acquisition of dendritic characteristics in the proximal axon ^3,72^. Such complications limit the dissection of the molecular mechanisms underlying gAnkG’s function. In contrast, targeted degradation technologies can acutely and effectively eliminate proteins within hours, thus providing a valuable temporal window for studying dynamic cellular processes especially for essential proteins whose chronic depletion causes cellular toxicity or organismal lethality.

We adopted a Trim21-based protein degradation system to achieve acute AIS degradation. Trim21, an E3 ubiquitin ligase, dimerizes to activate its ubiquitin ligase activity for proteasome-mediated degradation of its targets ^73^. In our design, a monomeric, inactive RING domain derived from Trim46 was fused to the N-terminus of the ABD within the EATT system for AIS targeting. The high-density clustering of gAnkG at the AIS is expected to promote dimerization of the targeted RING domain, thereby activating its E3 ligase activity, and enabling localized AIS protein degradation (schematic in Figure 7a). As shown in Figure 7b and 7c, hippocampal neurons transfected for 18 hours with a conventional ABD-fused Trim21 construct (i.e., a construct without the feedback expression control of the EATT system) completely lost their AIS structures as indicated by gAnkG staining. However, the majority of these neurons (about 75%) exhibited abnormal morphology, including membrane invaginations, irregularities, as well as neurite defects such as swelling, fragmentation, and aberrant processes that are unusually robust or slender, likely due to off-target Trim21 activities outside the AIS.

**Figure 7:**
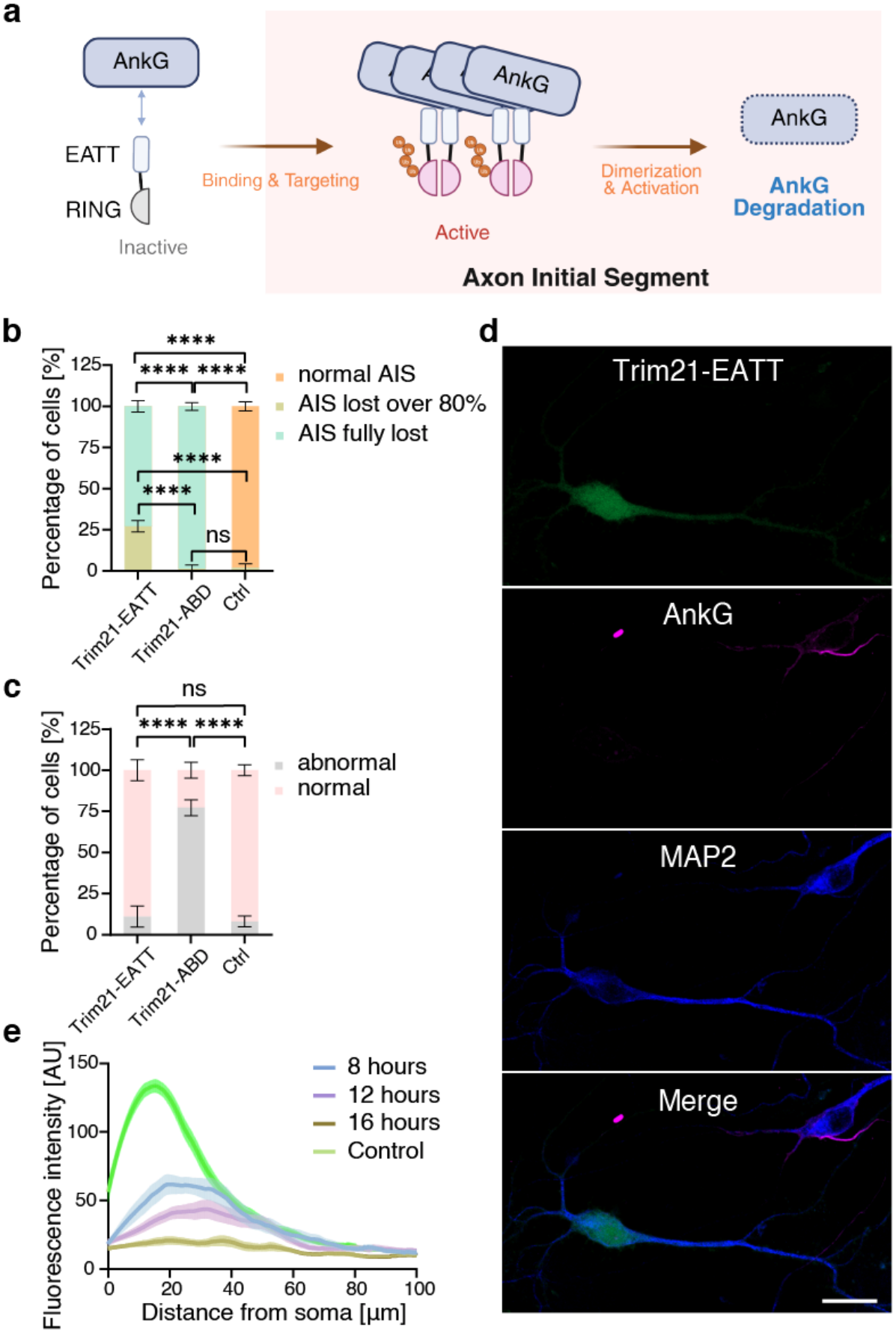
Acute degradation of gAnkG by the EATT system. **a:** Schematic diagram illustrating the principle of acute degradation of gAnkG at the AIS via the Trim21-based protein degradation pathway. **b:** AIS morphology was classified as fully degraded, obviously weaker, or normal. Bars represent the mean percentage of cells in each phenotype category, and error bars indicate SEM from three independent experiments. A total of 73 Trim21-EATT cells, 60 Trim21-ABD cells, and 55 control cells were analyzed. The complete three-category phenotype distributions were compared using generalized Cochran-Mantel-Haenszel tests stratified by independent experiment, followed by Holm correction for three prespecified pairwise comparisons. Trim21-EATT versus control, adjusted ****P<0.0001; Trim21-ABD versus control, adjusted ****P<0.0001; Trim21-EATT versus Trim21-ABD, adjusted ****P<0.0001. **c**: The percentage of cells displaying abnormal morphology was quantified in the Trim21-EATT, Trim21-ABD, and control groups. A total of 101 Trim21-EATT cells, 67 Trim21-ABD cells, and 99 control cells were analyzed. Statistical comparisons were performed using experiment-stratified Cochran-Mantel-Haenszel tests, followed by Holm correction for multiple pairwise comparisons. Trim21-EATT versus control, adjusted P=0.396; Trim21-ABD versus control, adjusted ****P<0.0001; Trim21-ABD versus Trim21-EATT, adjusted ****P<0.0001; n=3 independent experiments, mean ±SEM. **d:** Representative images of DIV12 neurons fixed 18 h after transfection with Trim21-EATT (green), showing loss of endogenous gAnkG (magenta). MAP2 is stained blue. AIS structures are absent in Trim21-EATT-transfected neurons. Scale bar: 20 μm. **e:** gAnkG immunostaining profiles along axons, showing rapid AIS degradation within hours of Trim21-EATT expression. n=15, 18, 15 and 29 neurons for 8h, 12h, 16h and control respectively, mean ± SEM.

To improve specificity, we employed the EATT2.0 system (Trim21-EATT) to restrict Trim21 expression at the AIS. This controlled and compartmentalized expression still induced efficient and near-complete loss of the AIS, as indicated by anti-pan-AnkG antibody staining (Figure 7b, 7d), while preserving overall normal neuronal morphology (Figure 7c). We next assessed the kinetics of AIS degradation by fixing neurons at 8, 12, and 16 hours after liposome-based transfection with Trim21-EATT and quantifying anti-pan AnkG signals at the AIS. As shown in Figure 7e, AnkG levels were significantly reduced by 8 hours, further diminished at 12 hours, and essentially absent at 16 hours. Taken together, these results demonstrate a highly efficient and specific tool for acute AIS degradation, providing a defined temporal window for investigating the role of AIS disassembly in neurodegeneration, axonal injury and other related pathological conditions.

## Discussion

The AIS functions as a pivotal hub for regulating neuronal electrical signaling and exhibits remarkable structural plasticity. This regulatory capacity is instrumental in determining action potential firing patterns, thereby positioning the AIS as an ideal locus for modulating overall neuronal excitability. Despite its critical roles, the full potential of the AIS as a neuronal activity regulatory switch has not been adequately exploited in both basic research and translational applications, largely due to the lack of tools capable of targeting to the AIS with high selectivity and robustness. The genetically encodable EATT system enables precise, endogenous-level AIS targeting with spatiotemporal and cell-type specificities. Its modular design allows easy adaptation for diverse applications, making the system a versatile platform for dissecting AIS physiology and associated pathophysiology.

The EATT builds on the previously reported FingR-based synaptic labeling system, which uses genetically encoded intrabodies (FingRs) targeting the synaptic scaffold proteins PSD95 or gephyrin combined with a CCR5- or IL2R-KRAB transcriptional feedback system to autoregulate tool expression and minimize nonspecific signals ^26^. While sharing similar design principle, our tool incorporates two key innovations to extend this strategy to the AIS. First, not all high-affinity gAnkG binders can achieve expected AIS targeting specificity, although the underlying mechanisms are unclear. The discovery of MCF2L-Spec1 as an effective AIS-targeting module providing the foundation for developing AIS-targeting tools. Second, the original transcriptional feedback system suffers from severe nuclear signal accumulation ^26,37–39^, which poses potential risks to nuclear processes such as gene transcriptions. Our design substantially alleviated this nuclear accumulation problem by increasing the copy numbers of the ZnF binding sequence.

The specificity of our EAAT system hinges on the identification of a highly stable, strong and specific gAnkG-binding protein domain containing only a single spectrin repeat. Structure-guided engineering further improved its binding affinity for gAnkG. The resolved structure of the gAnkG/Spec1 complex provides the molecular basis for further refinements when necessary. Notably, binding to gAnkG markedly enhances the structural stability of the spectrin repeat domain (Figure S6b). This induced stabilization likely extends the protein’s half-life in living cells and may confer additional specificity for the AIS labeling over other subcellular compartments. Additionally, the high conservation of the Spec1 binding sequence in gAnkG orthologs, spanning from cartilaginous fishes to humans, further indicates that our EAAT system can be used to study the AIS across diverse species.

In addition to labeling the AIS, the EATT developed in this study also labeled nodes of Ranvier, another type of micron-scale axonal domain that is enriched with gAnkG and structurally homologous to the AIS ^74^. EAATs were also detected in the somatic and proximal dendritic membranes of neurons, where gAnkG functions to stabilize somatodendritic GABAergic synapses ^75^, but the somatodendritic membrane signals of EAAT were substantially lower than at the AIS or nodes of Ranvier. Since cultured neurons lack nodes of Ranvier, the AIS constitutes the predominant site of EATT labeling, although residual signal persists in the somatic membrane, and faint nuclear signals were occasionally observed in a small subset of neurons.

Compared with the AnkG-GFP knock-in strategy ^69^, our method offers additional flexibility and broader applicability for studying AIS functions. The knock-in line minimally perturbs AIS organization and neuronal physiology, providing superior fidelity for long-term, minimally invasive AIS imaging both in vivo and in vitro, though it is limited to reporting AnkG localization and depends on Cre availability and transgenic breeding. Our EATT system, instead, enables delivery of essentially any desired proteins/organelles to the gAnkG-positive compartments for structural and functional investigations. When combined together, these two complementary and orthogonal methods offer great power for studying AIS biology.

## Acknowledgement

This work was supported by grants from the Major Program of Shenzhen Bay Laboratory (Grant No. S201101002), Shenzhen Medical Research Fund (E250200113), Shenzhen Talent Program (KQTD20210811090115021), Shenzhen Science and Technology Basic Research Program (JCYJ20220818100215033), Guangdong Innovative and Entrepreneurial Research Team Program (2021ZT09Y104), the Key Research and Development Program of Guangdong Province (Grant No. 2023B0303010001), the National Natural Science Foundation of China (Grant No. 32000874).

## Author contributions

K. C. and M. Z. conceptualized the project and designed the experiments. K. C., Y. L., F. L., M. S., Y. H., Q. T., W. C., W. Z. and H. M. performed all the experiments and analyzed the data. K. C. and M. Z. supervised the study. K. C., Y. L. and M. Z. wrote the original draft. All authors review and edited the manuscript. M. Z. and K. C. acquired the founding.

## Data, Materials, and Software Availability

All study data are included in the article and/or SI Appendix. All unique reagents generated in this study are available upon request from the corresponding author.

## Materials and Methods

The coding sequences of MCF2L were PCR-amplified from a mouse brain cDNA library. The coding sequences of AnkG constructs were derived from a full-length rat 480-kDa AnkG, kindly provided by Dr. Vann Bennett (Duke University). Details of other constructs and reagents, animals, protein sample preparation, biochemical assays, neuron culture and imaging and AIS quantification are described in the *SI Materials and Methods*.

