## Supplementary material for "Methods for precise targeting and modulation of the axon initial segment": Supplymentary information

**This PDF file includes:**

Supporting materials and methods
Figures S1 to S6
Tables S1
Legend for Movie S1
SI References

**Other supporting materials for this manuscript include the following:**

Movies S1

### Supporting materials and methods

#### Animal maintenance and welfare

The AnkG-GFP knock-in mouse strain was designed in-house and generated by Cyagen Co. Ltd (Suzhou, China) using CRISPR-Cas9 technology on a C57BL/6J genetic background. Heterozygous F1 mice were backcrossed with wild-type C57BL/6J mice for more than three generations before intercrossing to produce homozygous animals. All animal procedures were approved by the Institutional Animal Care and Use Committee of Shenzhen Bay Laboratory and conducted in accordance with relevant guidelines for laboratory animal welfare.

#### Plasmids and antibodies

The coding sequences of MCF2L were PCR-amplified from a mouse brain cDNA library. The MCF2L mutant constructs were generated by site-directed mutagenesis using the QuikChange II XL Site-Directed Mutagenesis Kit (Agilent Technologies) with the WT plasmid as the template. The coding sequences of AnkG constructs were derived from the full-length rat 480-kDa AnkG, a gift from Dr. Vann Bennett (Duke University). The human Synapsin-1 (hSyn) promoter was PCR amplified from pAAV-hSyn-GFP (Addgene #50465), the CamkIIα promoter was obtained from pAAV-CaMKIIa-hM3D(Gq)-mCherry (Addgene #50476), and the hDlx promoter was sourced from pAAV-hDlx-GqDREADD-dTomato-Fishell-4 (Addgene #83897). Additional plasmids, antibodies, and other related reagents are summarized in the Table S1.

#### Protein expression and purification

Recombinant proteins were expressed in BL21-CodonPlus (DE3) Escherichia coli grown in lysogeny broth (LB) medium. Protein expression was induced with 0.15 mM IPTG at 16°C for 20 hours. Cells were harvested and lysed, and the clarified supernatant was applied to $Ni^{2+}$-nitrilotriacetic acid agarose (Cytiva; 17531803) for initial purification via the N-terminal $His_6$-tag. The eluted protein was further purified by size-exclusion chromatography on a HiLoad 26/600 Superdex 200 pg column (Cytiva), equilibrated in buffer comprising 50 mM Tris-HCl (pH 7.8), 100 mM NaCl, 1 mM EDTA, and 1 mM DTT. The $His_6$-tag was cleaved by incubation with HRV-3C protease and removed by additional size-exclusion chromatography.

#### Isothermal titration calorimetry (ITC) assay

ITC measurements were performed on a MicroCal PEAQ-ITC calorimeter (Malvern Panalytical) following established procedures (1). The syringe was filled with ABD protein at 300 μM, which was titrated into the sample cell containing its binding partner AnkG at 30 μM. The reaction temperature was maintained at 25°C. Each experiment comprised 19 titration points, with a 120-second interval between injections, excluding the first data point from analysis. Data were analyzed and fitted using a one-site model within the manufacturer's software.

#### Size-exclusion chromatography with multi-angle light scattering (SEC-MALS) assay

SEC-MALS analysis was performed on a fast protein liquid chromatography (FPLC) system (AKTA pure L, Cytiva) coupled with a static light-scattering detector (miniDAWN; Wyatt) and a differential refractive index (dRI) detector (Optilab; Wyatt). Protein samples (100 μL) were loaded to a Superdex 200 Increase 10/300 GL column (Cytiva) pre-equilibrated with running buffer containing 50 mM Tris-HCl pH 7.8, 100 mM NaCl, 1 mM EDTA and 1 mM DTT. Data were collected and analyzed using Astra 8 software (Wyatt).

**Neuron culture, transfection and immunostaining**

Primary hippocampal or cortical neurons were dissected from neonatal brains (postnatal day 0-1 mice) and kept in cold HBSS/Hibernate buffer until processing. For plating, the tissue was enzymatically digested with trypsin and DNase I at 37 °C for 15 min, washed with HBSS, then mechanically dissociated by gentle trituration in plating medium (Neurobasal-A supplemented with 10% FBS, 2% B27, 1% GlutaMAX, and 1% Penicillin-Streptomycin). The cell suspension was filtered through a 70 μm cell strainer and then plated at a density of approximately 1,200,000 cells per dish onto 35-mm MatTek dishes, with a 10-mm diameter glass-bottom well pre-coated with poly-D-lysine and laminin (Sigma-Aldrich). Cultures were maintained at 37°C with 5% $CO_2$. After 6~12 hours, the medium was replaced with growth medium (Neurobasal-A supplemented with 1% FBS, 2% B27 and 1% GlutaMAX), designated as day in vitro 1 (DIV1). One-third of the medium was refreshed every 3~4 days until fixation.

Neurons were transfected at days in vitro 4 (DIV4, unless otherwise specified) using Lipofectamine 2000 (Thermo Fisher). For immunostaining, neurons were fixed at room temperature for 15 minutes with 4% (wt/vol) paraformaldehyde (PFA) and 4% (wt/vol) sucrose in phosphate-buffered saline (PBS) (For pan-NaV immunostaining, 0.5% PFA for 20 min was used instead). After fixation, neurons were washed with PBS, permeabilized with 0.25% Triton X-100 in PBS for 5 minutes, and blocked with 5% bovine serum albumin (BSA) in PBS for 1 hour. Primary antibodies (see antibody list and their dilution ratios in the Table S1) diluted in blocking buffer, were applied overnight at 4°C. On the following day, neurons were rinsed with PBS and incubated with the appropriate fluorescent secondary antibodies prepared in blocking buffer for 1 hour at room temperature. Finally, neurons were washed twice with PBS and stored in PBS for imaging.

**Confocal microscopy and live cell imaging**

Imaging of fixed neurons was performed on a Zeiss LSM980 laser scanning confocal microscope (Axio Observer inverted platform) equipped with an LD C-Apochromat 40×/1.1 W Korr UV VIS IR water-immersion objective. Multi-channel fluorescence images were acquired sequentially using a multitrack configuration with frame-by-frame switching to prevent spectral bleed-through. Each optical section was averaged over 2 frames (pixel dwell time: 0.51 μs) to improve the signal-to-noise ratio. Three-dimensional Z-stacks covering the entire neuronal thickness were acquired at a resolution of 2048×2048 pixels, with a pinhole size of 1.0~2.0 Airy Units (AU) and a Z-step size of ~1.0 μm. Laser power and detector gain were optimized for each independent experiment to prevent signal saturation, and identical parameters were maintained across all experimental groups.

Live cell imaging was performed on the same confocal system equipped with an on-stage incubation chamber maintaining stable temperature and $CO_2$ levels. Neurons were cultured in phenol red-free medium to minimize background fluorescence. Automated time-lapse imaging was performed at DIV13 by combining tile-scan and Z-stack modalities across approximately 30 pre-selected random positions. Following KCl addition to induce plasticity, the positions were scanned continuously at 15~30 min intervals. At each position, Z-stacks (512×512 pixels) covering the full thickness of all selected neurons were acquired with a pinhole size of 2.3 AU and a Z-step size of ~1.0 μm. Each optical section was averaged over 2 frames (pixel dwell time: 1.02 μs). Imaging continued for 30~48 hours, with real-time Definite Focus used to counteract axial drift.

**Image processing and AIS analysis**

Raw Z-stacks were processed using ZEISS ZEN (Blue edition) software to generate maximum intensity projections (MIPs). The MIP images were exported as PNG files for analysis or figure preparation. For better visualization, some images underwent brightness adjustments; identical processing parameters were applied to all images within each experimental group to ensure comparability. No nonlinear adjustments or other image manipulations were performed.

The AIS was identified using ABD-GFP fluorescence without immunostaining or by immunolabeling endogenous AnkG. All dishes were processed under identical labeling conditions. Morphometric analysis was performed using Fiji/ImageJ: fluorescence intensity profiles were obtained by drawing line scans (line width, 6 pixels) along the axon, starting from the soma boundary and extending for about 100-200 μm. For AISs originating from dendrites, the line scans started from the boundary of the dendritic shaft instead. The resulting profiles were analyzed using a custom-written Python script implementing Savitzky-Golay smoothing with a second-order polynomial. For baseline estimation, the terminal 10% of each valid, smoothed profile was defined as the putative steady-state region. The lowest and highest 10% of values within this region were excluded, and the mean of the remaining values was used as the curve-specific baseline.

After baseline subtraction, the peak fluorescence intensity was determined for each smoothed profile. A threshold corresponding to 33% of the peak intensity was used to identify the AIS boundaries, including the start and end positions. Intersections between the smoothed fluorescence profile and the threshold were calculated by linear interpolation. The AIS starting point was defined as the x-coordinate of the first intersection; if the smoothed fluorescence profile had only one intersection with the threshold, the AIS starting point was defined as x=0. The soma-AIS gap was defined as the distance between the soma (or dendrite) boundary (x-coordinate of zero) and the x-coordinate of the first intersection (if the smoothed fluorescence profile has only one intersection with the threshold, the soma-AIS gap distance is zero), and AIS length was defined as the largest distance between two consecutive threshold intersections. Each automatically processed AIS profile was manually inspected to ensure consistency with the corresponding original image and raw line-plot curve.

For AIS fluorescence intensity quantification (Figure 2g), a contiguous 10 μm segment exhibiting the highest mean fluorescence intensity along each AIS fluorescence profile was identified using a sliding-window approach. The mean fluorescence intensity within this selected segment was then calculated and used as a representative measure of the overall AIS fluorescence intensity.

**Virus preparation, mouse brain injection and immunohistochemistry**

HEK293T cells were transfected at 70-80% confluency using a triple-plasmid cocktail at a molar ratio of 1:1:1, comprising the plasmid of interest, the pAAV-helper (Cell Biolabs, Inc.), and the AAV2/9 capsid plasmid (Addgene #112865). Transfection was performed with 1 mg/mL PEI-MAX (Polysciences) at a PEI-to-DNA ratio of 4:1 (w:w). The PEI and plasmid DNA were diluted in DMEM, incubated at room temperature for 15 minutes, and then gently added dropwise to the cells.

After a 3-day transfection period, the medium was removed, and cells were rinsed with gradient buffer (100 mM Tris-HCl, pH 7.5, 150 mM NaCl, 10 mM $MgCl_2$) before being gently scraped and collected. The harvested cells were lysed by four freeze-thaw cycles, treated with Omnipotent Nuclease (7seabiotech, China) at 37°C for 60 minutes, and centrifuged at 3,500× g for 15 minutes at 10°C.

For *in vitro* infection, the crude AAV supernatant was filtered through a 0.45 μm PES filter and concentrated using an Amicon Ultra Centrifugal Filter (30 kDa MWCO). For higher purity, the lysate was subjected to iodixanol gradient centrifugation (15%, 25%, 40%, and 58%) in a Beckman Quick-seal ultracentrifuge tube at 48,000 rpm for 2 hours and 10 minutes at 16°C. The AAV fraction extracted from the 40% to 58% interface was concentrated using an Amicon Ultra Centrifugal Filter (100 kDa MWCO). The purified AAV was aliquoted and stored at -80°C for long-term preservation.

AAV injection into the cortex and striatum was conducted as follow: Adult C57BL/6J mice were anesthetized with ketamine (80 mg/kg) and xylazine (10 mg/kg). A craniotomy was performed to expose the cortex. For cotical injection, AAVs ($1\times10^{13}$ GC/mL) were injected at two coordinates (anteroposterior/mediolateral/dorsoventral): 0, 2, -0.6; and 1, 1.5, -0.6. For striatal injection, AAVs ($1\times10^{13}$ GC/mL) were injected at two coordinates: 0, 2, -2.5; and 1, 1.5, -2.5. A total of 500 nL AAV was infused into each site at a rate of 100 nL/min. The skin was sutured after injection. The animals were placed on a heating pad until they were fully awake.

For immunohistochemistry, mice were euthanized by transcardiac perfusion, and the brains were immediately dissected and post-fixed overnight in 4% (wt/vol) paraformaldehyde at 4 °C. Fixed brains were then cryoprotected in 30% (wt/vol) sucrose in PBS until they sank completely. Brains were embedded in OCT compound (Tissue-Tek), frozen on dry ice, and stored at −80 °C until sectioning. Frozen sections were cut to a thickness of 15 μm using a Leica CM3050S cryostat and collected in PBS for free-floating staining or mounted directly onto gelatin-coated slides. Prior to staining, sections were washed three times for 5 min each in PBS to remove residual OCT compound. Antigen retrieval was avoided to preserve GFP signals. Sections were blocked with blocking buffer for 1 h at room temperature. The slides were then incubated overnight at 4 °C with primary antibodies diluted in blocking buffer. Following extensive washes with PBS, sections were incubated with the appropriate secondary antibodies in blocking buffer for 1 h at room temperature, protected from light. After a final wash, slides were mounted with ProLong™ Antifade reagent (Thermo Fisher) and coverslipped.

**Quantification and statistical analysis**

Data are presented as mean ± SEM. Statistical analyses other than those for Figures 7b and 7c were performed using GraphPad Prism 8.4.3. Data distributions were assessed for normality using the Shapiro-Wilk or Kolmogorov-Smirnov test, as appropriate. Comparisons between two groups were performed using a two-tailed Student's t-test for normally distributed data or a two-tailed Mann-Whitney U test for nonnormally distributed data. For Figures 7b and 7c, phenotype counts were analyzed using Cochran-Mantel-Haenszel tests stratified by independent experiment, with generalized tests used for the three-category outcome in Figure 7b and 2×2 tests used for the binary outcome in Figure 7c. Two-sided P values from three prespecified pairwise comparisons were adjusted using the Holm method, and analyses were performed in Python 3.12.0 using NumPy, SciPy, and statsmodels. Statistical significance was defined as $P < 0.05$, with the following notations: ns, $P \geqslant 0.05$; *$P < 0.05$; **$P < 0.01$; ***$P < 0.001$; ****$P < 0.0001$.

### Figures

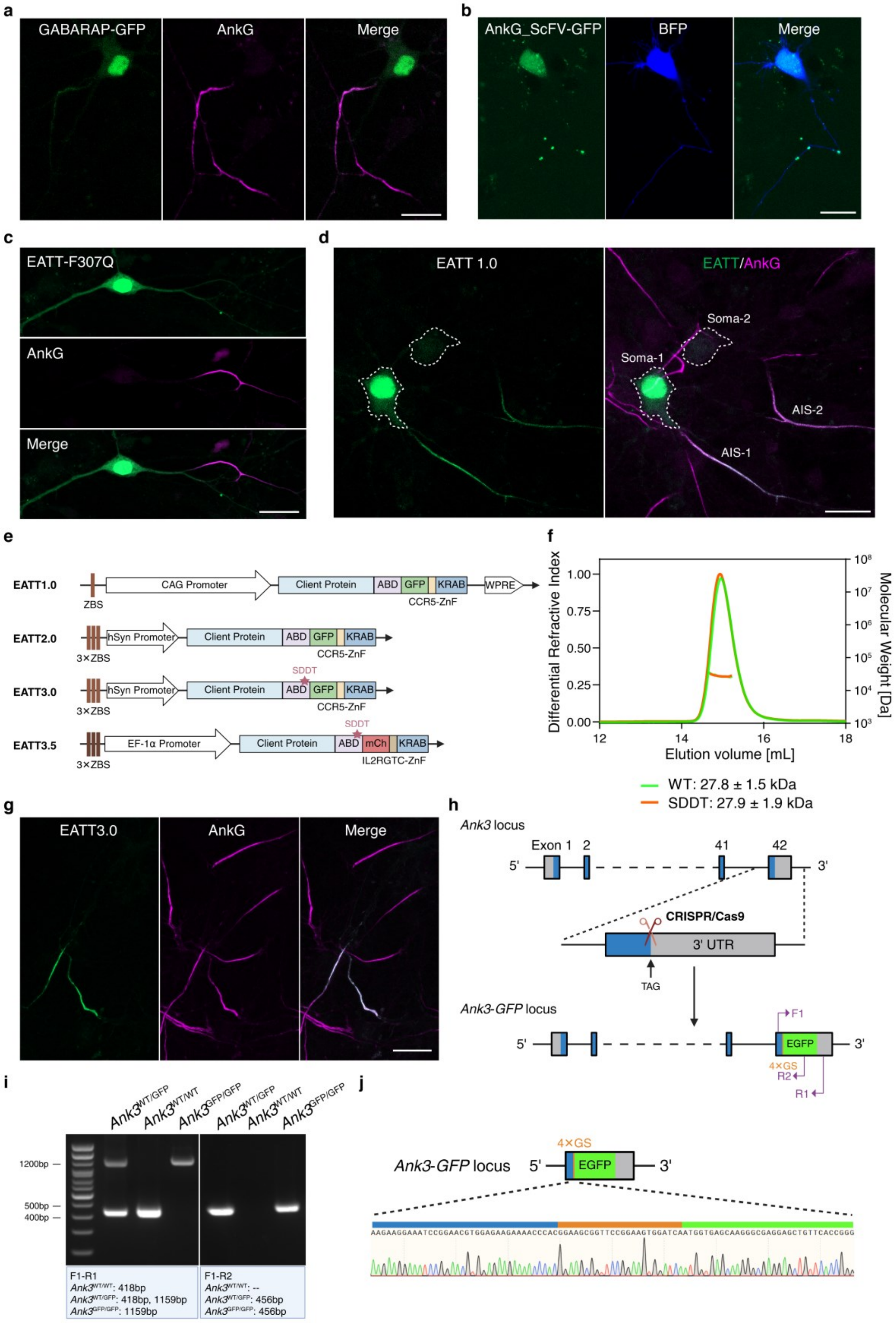

a
GABARAP-GFP
AnkG
Merge
b
AnkG_ScFV-GFP
BFP
Merge
c
EATT-F307Q
AnkG
Merge
d
EATT 1.0
EATT/AnkG
Soma-1
Soma-2
AIS-1
AIS-2
e
EATT1.0
ZBS
CAG Promoter
Client Protein
ABD
GFP
KRAB
WPRE
CCR5-ZnF
EATT2.0
3×ZBS
hSyn Promoter
EATT3.0
SDDT
EATT3.5
EF-1α Promoter
mCh
IL2RGTC-ZnF
f
Differential Refractive Index
Molecular Weight [Da]
Elution volume [mL]
WT: 27.8 ± 1.5 kDa
SDDT: 27.9 ± 1.9 kDa
g
EATT3.0
AnkG
Merge
h
Ank3 locus
Exon 1
2
41
42
5'
3'
CRISPR/Cas9
3' UTR
TAG
Ank3-GFP locus
F1
EGFP
4×GS
R2
R1
i
Ank3WT/GFP
Ank3WT/WT
Ank3GFP/GFP
1200bp
500bp
400bp
F1-R1
Ank3WT/WT: 418bp
Ank3WT/GFP: 418bp, 1159bp
Ank3GFP/GFP: 1159bp
F1-R2
Ank3WT/WT: --
Ank3WT/GFP: 456bp
Ank3GFP/GFP: 456bp
j
Ank3-GFP locus
AAGAAGGAAATCCGGAACGTGGAGAAGAAAACCCACGGAAGCGGTTCCGGAAGTGGATCAATGGTGAGCAAGGGCGAGGAGCTGTTCACCGGG

**Fig. S1. Design and validation of the EATT system.**

**a-b:** Images demonstrating that GABARAP and gAnkG_scFv failed to label the AIS in cultured hippocampal neurons. Scale bar: 20 μm.

**c:** A single point mutation (F307Q) in the gAnkG-binding domain (ABD, specifically the first spectrin repeat) of MCF2L within the EATT construct abolishes the targeting of ABD-GFP to the AIS. Scale bar: 20 μm.

**d:** Example of EATT1.0 transfected neurons showing nuclear ABD-GFP accumulation. Scale bar: 20 μm.

**e:** Schematic diagram illustrating the plasmid constructs of various EATT versions.

**f:** Multi-angle static light scattering results showing that the SDDT mutant of the MCF2L ABD elutes at the same volume as the wild-type (WT) protein during size-exclusion chromatography and exhibits the same molecular weight.

**g:** EATT3.0 achieved near-exclusive AIS localization. Magenta: immunostained gAnkG marking the AIS. Green: ABD-GFP fluorescence signal from EATT3.0. Scale bar: 20 μm.

**h:** Design strategy for generating AnkG-GFP knock-in mice by inserting an EGFP tag at the C-terminus of ankyrin-G.

**i-j:** Genotyping PCR and sequencing results confirming the homozygous AnkG-GFP knock-in mice.

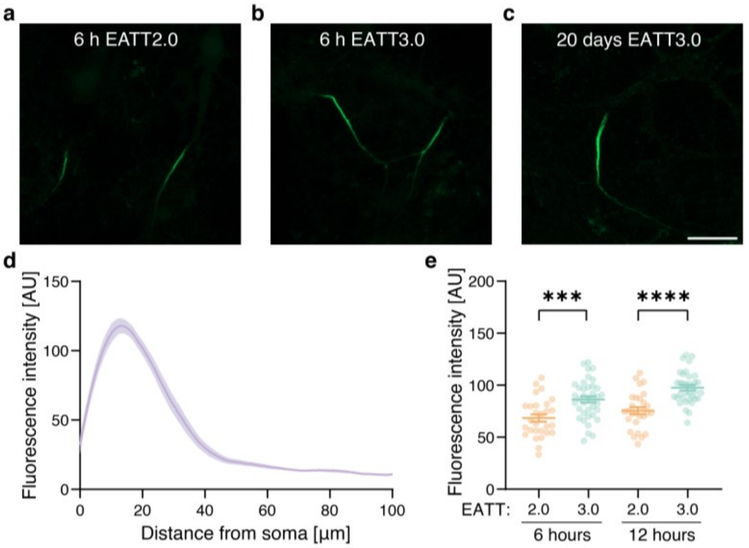

**Fig. S2. Time window of the EATT to label AIS.**

**a-b:** Representative images of hippocampal neurons transfected with EATT2.0 or EATT3.0 at DIV8 and imaged for GFP signal 6 h post-transfection without immunostaining. Images were acquired under identical settings. Scale bar: 20 μm.

**c-d:** EATT specifically marks the AIS in a high signal-to-noise ratio even after 20 days' expression in culture. Representative image and intensity profiles of neurons transfected with EATT3.0 at DIV4 and imaged for GFP signal at DIV24. Images were taken under the same microscope settings as in panel a and b. Curves are smoothened.

**e:** Cultured hippocampal neurons transfected with EATT2.0 or EATT3.0 plasmids at DIV8, and GFP signal was quantified without immunostaining at 6 hours or 12 hours post-transfection. Each data point represents one AIS. Student's t-test, n>28 per group, mean $\pm$ SEM.

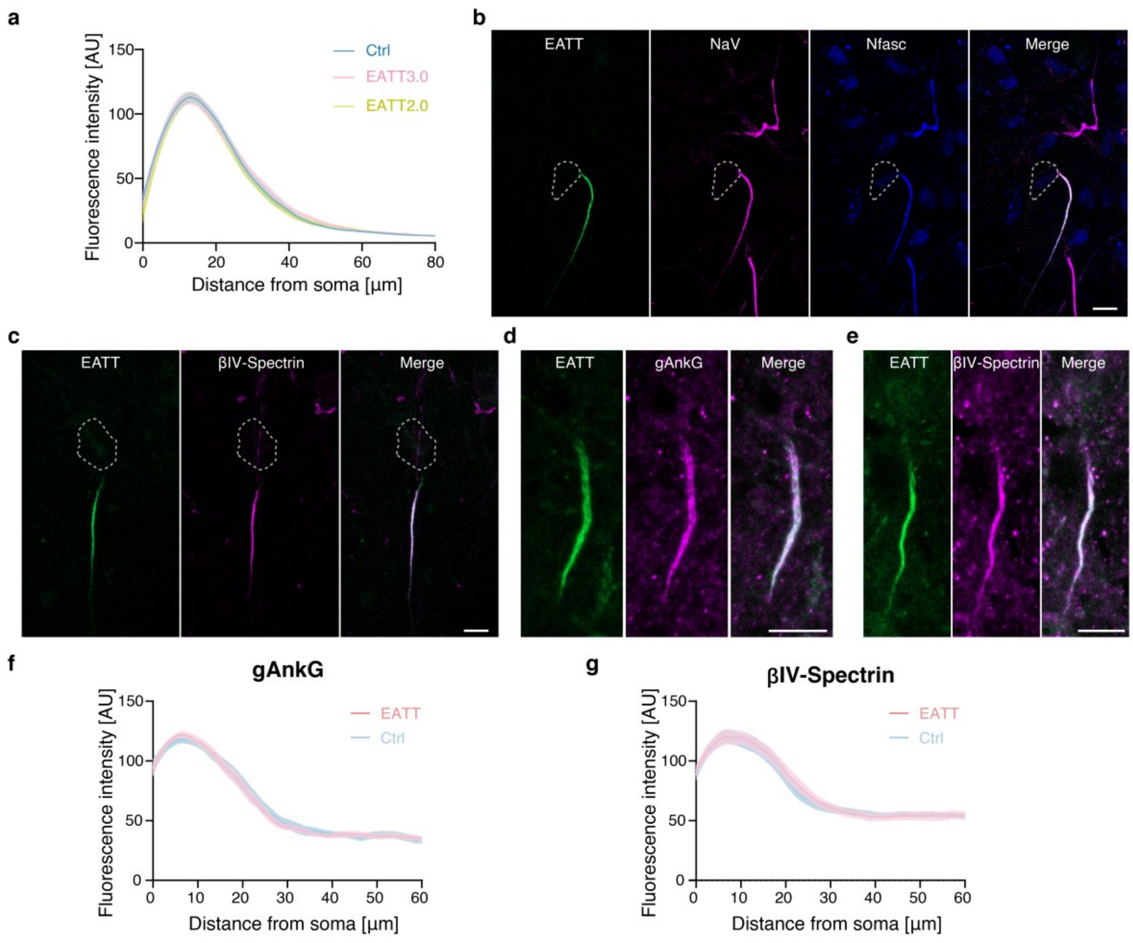

**Fig. S3. EATT does not change AIS structures *in vitro* and *in vivo*.**

**a:** Line-plot profiles of AISs from EATT2.0 or EATT3.0 transfected neurons or non-transfected control neurons showing that EATT2.0 or EATT3.0 does not alter AIS morphology. Cultured hippocampal neurons were transfected at DIV4 and imaged at DIV13. Curves are smoothened for better presentation; n=38 for Ctrl; n=31 for EATT2.0; n=31 for EATT3.0.

**b-c:** Expended optical views corresponding to Figure 3c and 3f. Scale bar: 10 µm.

**d-g:** Neurons from frozen mouse cortical slices infected with AAV9-packaged EATT2.0 show perfect colocalization with immunostained gAnkG or βIV-spectrin. Scale bar: 10 µm.

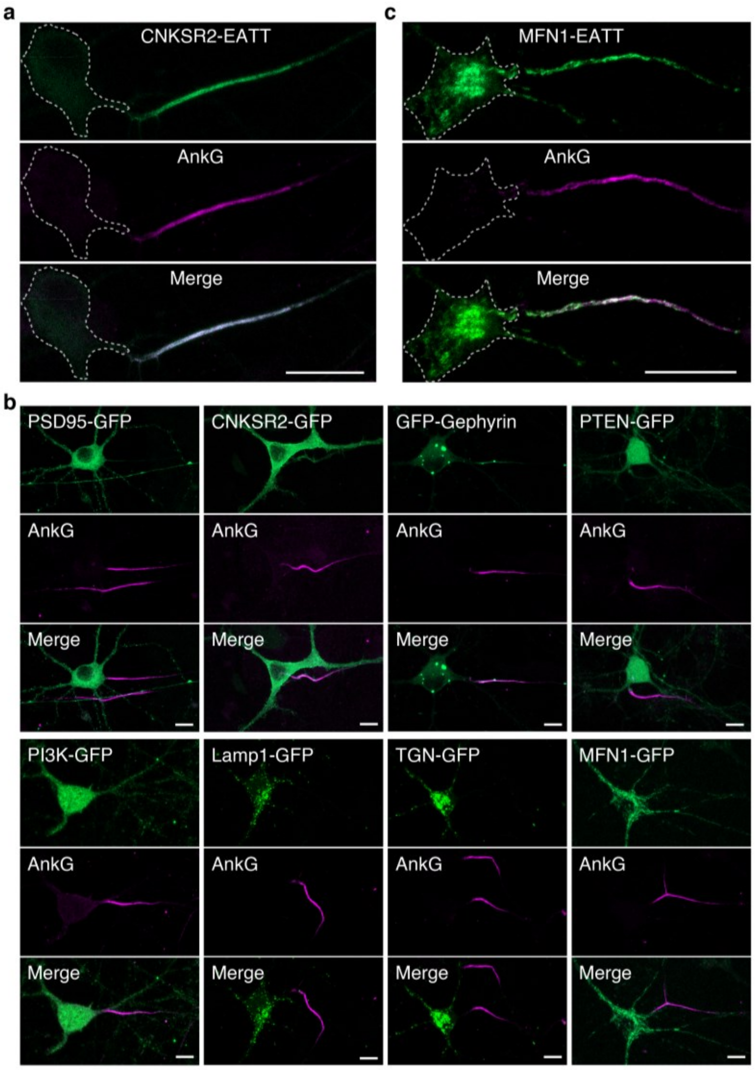

**Fig. S4. Expanded optical views of neurons in Figure 3.**

**a:** Representative images showing CNKSR2 was enriched at the AIS by CNKSR2-EATT. Endogenous gAnkG was immunostained with a mouse anti-AnkG antibody. Scale bar: 20 μm.

**b:** Expended optical views corresponding to neurons in Figure 3h-m. Scale bar: 10 μm.

**c:** Representative images showing MFN1 marked mitochondria was enriched at the AIS by MFN1-EATT. Endogenous gAnkG was immunostained with a mouse anti-AnkG antibody. Scale bar: 20 μm.

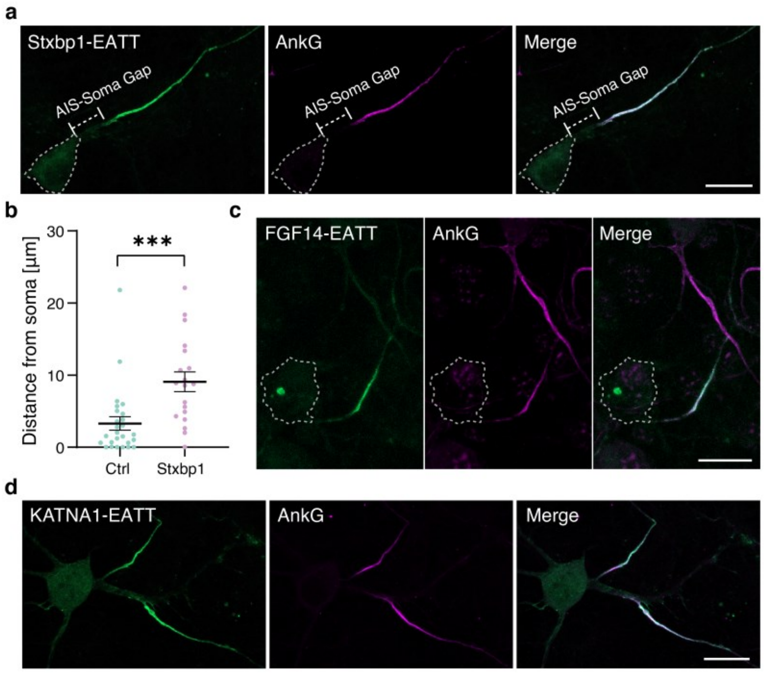

**Fig. S5. Additional examples of modulating AIS structure using the EATT system and newly characterized modulators.**

**a-b:** Quantification showing that Stxbp1-EATT significantly increased the AIS-soma gap distance. Scale bar: 10 µm. Mann Whitney test, n=19 (Stxbp1-EATT) and 26 (Ctrl), mean ± SEM.

**c-d:** Representative images of hippocampal neuron AIS targeted by FGF14-EATT (**c**) or KATNA1-EATT (**d**). Endogenous gAnkG was immunostained with a mouse anti-AnkG antibody. Scale bar: 20 µm.

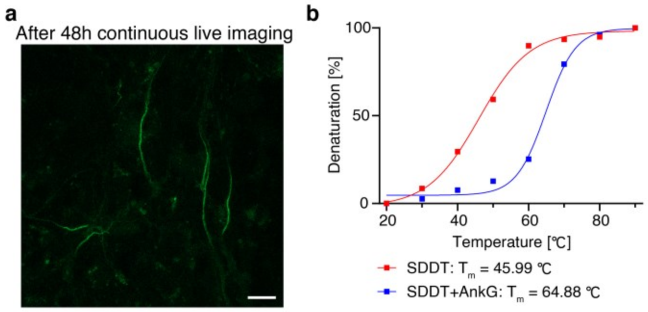

**Fig. S6. Long-term monitoring of AIS structure using EATT in living neurons.**

**a:** Cultured hippocampal neuron AIS labeled by ABD-Baojin (Green) using EATT3.0, showing a bright fluorescent signal after 48 hours of continuous live-cell imaging under a confocal microscope. Scale bar: 10 µm.

**b:** Absorbance changes at a wavelength of 222 nm in the circular dichroism (CD) spectra during heat-induced denaturation experiments of the ABD SDDT mutant, either alone or in complex with the gAnkG fragment. Binding to the gAnkG fragment greatly increased the stability of the ABD.

**Tables**

**Table S1. Supporting resource table.**

| **REAGENT or RESOURCE** | **SOURCE** | **IDENTIFIER** |
|---|---|---|
| **Antibodies** | | |
| Mouse monoclonal anti-Ankyrin-G (1:500) | NeuroMab | Cat#75-146; RRID: AB_10673030 |
| Chicken polyclonal anti-Ankyrin-G (1:1500) | Synaptic Systems | Cat#386006; RRID: AB_2814936 |
| Rabbit polyclonal anti-giant Ankyrin-G (1:1000) | Vann Bennett (Duke University) | N/A |
| Mouse monoclonal anti-sodium channel (1:100) | Sigma-Aldrich | Cat#S8809; RRID: AB_477552 |
| Rabbit monoclonal anti-Neurofascin 186 (1:100) | Cell Signaling Technology | Cat#15034S; RRID: AB_2773024 |
| Rabbit polyclonal anti-β4 spectrin (1:1500) | Vann Bennett (Duke University) | N/A |
| Mouse monoclonal anti-MAP2 (1:2000) | Sigma-Aldrich | Cat#M4403; RRID: AB_477193 |
| Rabbit monoclonal anti-CaMKII (1:400) | Beyotime | Cat#AF1639; RRID: AB_3711723 |
| Mouse monoclonal anti-GAD67 (1:500) | Sigma-Aldrich | Cat# MAB5406; RRID:AB_2278725 |
| Guinea pig monoclonal anti Parvalbumin (PV) (1:500) | Synaptic Systems | Cat#195308; RRID:AB_2927389 |
| **Bacterial and virus strains** | | |
| Escherichia coli BL21-CodonPlus(DE3)-RIL cells | Agilent Technologies | Cat#230245 |
| AAV-CaMKIIα-ABD-GFP-CCR5TC-KRAB-WPRE | This paper | N/A |
| AAV-hDlx-ABD-GFP | This paper | N/A |
| AAV-hSyn-ABD-mBaoJin-CCR5TC-KRAB-WPRE | This paper | N/A |
| **Chemicals, peptides and recombinant proteins** | | |
| Recombinant protein: MCF2L_ABD-WT (aa 196R-333V; NCBI: NM_178076.4) | This paper | N/A |
| Recombinant protein: MCF2L_ABD-SDDT (S280ESLE, D284Q, D287R, T291Q) | This paper | N/A |
| Recombinant protein: AnkG (aa 2482P-2615S) | This paper | N/A |
| XE991 | MedChemExpress | Cat#HY-108577A |
| Neurobasal™-A Medium | minus phenol red | Cat#12349015 |
| B27 | Gibco | Cat#17504044 |
| GlutaMAX™ Supplement | Gibco | Cat#35050061 |
| Penicillin-Streptomycin | Gibco | Cat#15140122 |
| DNaseI | Sigma-Aldrich | Cat#DN25 |
| Trypsin | ThermoFisher | Cat#15090046 |
| Fetal Bovine Serum (FBS) | Gibco | Cat#A5669701 |
| HBSS | Gibco | Cat#14170112 |

| | | |
|---|---|---|
| PEI | Polysciences | Cat#24765 |
| **Critical commercial assays** | | |
| Lipofectamine2000 transfection kit | Invitrogen | Cat#11668019 |
| **Experimental models: Organisms/strains** | | |
| Mice: C57BL/6J | Cyagen | N/A |
| Mice: *Ank3*$^{GFP}$ | Cyagen | Custom order |
| **Oligonucleotides** | | |
| *Ank3*$^{GFP}$ Forward primer (F1): 5'-CTCGTTATGCTGTCATTACCTTGA-3' | Sangon Biotech | Custom order |
| *Ank3*$^{GFP}$ Reverse primer (R1): 5'-TGGTGCAGATGAACTTCAGGGTC-3' | Sangon Biotech | Custom order |
| *Ank3*$^{GFP}$ Reverse primer (R2): 5'-CAGTGAGATTTCAGGAAAGAAGG-3' | Sangon Biotech | Custom order |
| **Recombinant DNA** | | |
| Plasmid: AnkG_scFv-GFP-CCR5TC-KRAB-WPRE | This paper | N/A |
| Plasmid: GABARAP-GFP-CCR5TC-KRAB-WPRE | This paper | N/A |
| Plasmid: EATT1.0 | This paper | N/A |
| Plasmid: EATT1.0-F307Q | This paper | N/A |
| Plasmid: EATT2.0 | This paper | N/A |
| Plasmid: EATT3.0 | This paper | N/A |
| Plasmid: EATT3.5 | This paper | N/A |
| Plasmid: PSD95-EATT | This paper | N/A |
| Plasmid: CNKSR2-EATT | This paper | N/A |
| Plasmid: Gephyrin-EATT | This paper | N/A |
| Plasmid: PTEN-EATT | This paper | N/A |
| Plasmid: PI3K-EATT | This paper | N/A |
| Plasmid: Lamp1-EATT | This paper | N/A |
| Plasmid: TGN38-EATT | This paper | N/A |
| Plasmid: MFN-EATT | This paper | N/A |
| Plasmid: λPPase-EATT | This paper | N/A |
| Plasmid: CDK5-EATT | This paper | N/A |
| Plasmid: 6×KBS-EATT | This paper | N/A |
| Plasmid: Stxbp1-EATT | This paper | N/A |
| Plasmid: FGF14-EATT | This paper | N/A |
| Plasmid: Trim46-EATT | This paper | N/A |
| Plasmid: KATNA1-EATT | This paper | N/A |
| Plasmid: Trim21-EATT | This paper | N/A |
| 3x $ZBS_{WT}$ | This paper | gtcatcctcatcactgtcatcctcatcagagtcatcctcatc |
| 6x $ZBS_{mut}$ + 1x $ZBS_{WT}$ | This paper | gtcatcgtcatcgtcgacattgattattgactgcagacatccacatctgagtcaacctcaacttagtcaaccacatcttggtcatccacatcactgtcatcgtcatcagagtcatcctcatc |
| **Software and algorithms** | | |
| Origin7.0 | OriginLab | http://www.originlab.com/ |
| PyMOL 1.5.x | PyMOL | http://www.pymol.org/ |

| | | |
|---|---|---|
| ImageJ-FIJI | NIH | https://imagej.net/ij/ |
| GraphPad Prism | GraphPad Software Inc | http://www.graphpad.com/scientificsoftware/prism/ |

**Movie S1** (separate file). Real-time monitoring of the interneuron AIS plasticity by EATT. Living imaging of AIS plasticity induced by KCl. Images were taken every 20 min; video playback at 10 frames per second.
